# Acidity promotes tumor progression by altering macrophage phenotype in prostate cancer

**DOI:** 10.1101/478420

**Authors:** Asmaa El-Kenawi, Chandler Gatenbee, Mark Robertson-Tessi, Rafael Bravo, Jasreman Dhillon, Yoganand Balagurunathan, Anders Berglund, Naveen Visvakarma, Arig Ibrahim-Hashim, Jung Choi, Kimberly Luddy, Robert Gatenby, Shari Pilon-Thomas, Alexander Anderson, Brian Ruffell, Robert Gillies

## Abstract

Tumors rapidly ferment glucose to lactic acid even in the presence of oxygen, and coupling high glycolysis with poor perfusion leads to extracellular acidification. Here we demonstrate that acidity, independent from lactate, augments the pro-tumor phenotype of macrophages. We used zwitterionic buffers to show that activating macrophages at pH 6.8 *in vitro* enhanced an IL-4-driven phenotype as measured by gene expression, cytokine profiling, and functional assays. These results were recapitulated *in vivo* wherein neutralizing intratumoral acidity reduced the pro-tumor phenotype of macrophages, while also decreasing tumor incidence and invasion in the TRAMP model of prostate cancer. These results were recapitulated using an *in silico* mathematical model that simulate macrophage responses to environmental signals. By turning off acid-induced cellular responses, our *in silico* mathematical modeling shows that acid-resistant macrophages can limit tumor progression. In summary, this study suggests that tumor acidity contributes to prostate carcinogenesis by altering the state of macrophage activation.

## Background

Cancer initiation and progression involves complex cellular interactions of pre-malignant/malignant cells with immune, stromal cells and blood vessels. Levels of tissue oxygen, metabolic byproducts, nutrients, and hormones modulate these cellular interactions that, in turn, can regulate tumor progression (1). One important property of malignant cells is that they preferentially metabolize glucose into lactate even in the presence of oxygen – known as aerobic glycolysis or the “Warburg Effect” – which confers on them a growth advantage (2). Coupling elevated glycolysis with poor tumor perfusion leads to increased pericellular accumulation of organic acids (e.g. lactic acid) and reduced pH in extracellular spaces (3). Low pH induces the activity of proteolytic enzymes and can be toxic to surrounding stromal cells, leading to tissue remodeling and local invasion (4, 5). It is also known to inhibit T cell mediated immune surveillance (6), but the effect of tumor acidosis on the myeloid compartment within tumor is less well studied.

Tumors are infiltrated by populations of myeloid cells that regulate tumorigenesis through their ability to mediate immunosuppression, matrix remodeling, angiogenesis, local invasion and metastasis (7, 8). In particular, infiltration by macrophages can promote tumor progression and poor outcome in solid malignancies when their presence is associated with a tumor-promoting phenotype reminiscent of IL-4-driven activation (9). The pro-tumor phenotype of these tumor-associated macrophages (TAMs) can be affected by several aspects of the tumor microenvironment (10). These include cytokines and antibodies produced by lymphocytes and tumor-derived cytokines/chemokines that promote macrophage infiltration and polarization (11-13). Abnormal metabolic factors can also aggravate the phenotype of these cells. For example, hypoxia augments the immunosuppressive ability of TAMs (14), while lactic acid induces tissue remodeling though expression of VEGF and arginase I (15). Whether acidic pH, as an independent entity from lactate (16), alters macrophage polarization within tumors is not clear; hence, we sought to investigate the impact of tumor acidosis on the phenotypic characteristics of macrophages *in vitro* using zwitterionic organic buffering agents. We then used a series of mouse models to correlate tumor progression with macrophage infiltration and to delineate the role of acidity in prostate cancer. We then reiterate our findings using an agent-based mathematical model that simulate how pH affects the ability of macrophages to control tumor growth.

## Materials and Methods

### Animal models

All mice were maintained in accordance with Institutional Animal Care and Use Committee (IACUC) standards followed by the Moffitt Cancer Research Center (Tampa, Florida). All animals and cell lines were male or male-derived, respectively, since this study is mainly investigating prostate cancer. For bone marrow isolation C57BL/6N (C57BL/6NHsd) 8-12 weeks male mice were purchased from Envigo. For the subcutaneous prostate cancer model, mice randomly assigned to experimental groups then provided with 200 mM bicarbonate in their drinking water 4 days prior to subcutaneous injections with 5×10^5^ TRAMP-C2 cells (17). Tumor growth was evaluated weekly by measurement of two perpendicular diameters of tumors with a digital caliper. Individual tumor volumes were calculated as volume = [π/6× (width)^2^ × length]. On the 35^th^-42^nd^ day post injection, solid tumors were harvested and processed for flow cytometric analysis and immunohistochemistry. Male Transgenic Adenocarcinoma of the Mouse Prostate (TRAMP) mice were obtained from The Jackson Laboratory. Male TRAMP spontaneously develops autochthonous prostate tumors following the onset of puberty due to the expression of the oncoprotein, SV40 T antigen (TAg) under transcriptional control of the rat probasin promoter (18).

### Cell lines

Male derived murine TRAMP-C2 and TRAMP-C3 prostate cancer cell lines were purchased from ATCC, maintained and cultured according to their suggested protocols.

### Macrophage isolation, activation and cell culture protocols

Bone marrow derived macrophages (BMDMs) were generated as described previously (19, 20). In brief, bone marrow was flushed from femurs and tibias of male C57BL/6N mice and cultured for 6-7 days in complete macrophage medium (Dulbecco modified Eagle's minimal essential medium (DMEM) supplemented with 10% fetal calf serum (FCS), 2% penicillin/streptomycin-glutamine) and 20 ng/ml M-CSF at 37°C. Pro-inflammatory macrophages were induced by exposing BMDMs to 50 ng/ml IFN-γ and 10 ng/ml LPS in complete macrophage medium. Anti-inflammatory macrophages were stimulated by exposure to 10 ng/ml IL-4 in complete macrophage medium (19, 21). Control macrophages (M0) were cultured for the same period in medium alone. Prostate cancer-associated macrophages were induced by incubating BMDMs with 30% 72 hr-conditioned medium from either TRAMP-C2 or TRAMP-C3 cell lines. To detect the effect of tumor microenvironmental acidity, macrophages were induced according the previous protocol but with further supplementation of media with the zwitterionic organic buffers PIPES and HEPES (25 mM each) and adjustment of the pH to either 7.4 or 6.8 (22).

### Antibodies, chemicals and kits

Recombinant mouse IFN-γ, M-CSF and interleukin (IL)-4 were obtained from R&D Systems. Sources of conjugated antibodies were as follows: iNOS-Alexa Fluor 488 (eBioscience), CD206-Alexa Fluor 647 (AbD Serotec), CD45-APC and MHCII-BV21 (BD Biosciences), F4/80-PE, Ly6C-APC/Cy7 and CD11b-PE/Cy7 (BioLegend). Sources of unconjugated antibodies were as follows: anti-MRC1 (CD206) and anti-iNOS (Abcam). Sources of chemicals were as follows: Rhodamine Phalloidin (Life Technologies). Griess reagent (Promega) was used to measure nitrite level. Click-iT EdU pacific blue flow cytometry assay kit (Life Technologies) was used to measure cell proliferation. Proteome profiler mouse cytokine array panel A or XL Cytokine Array ARY028 (R&D Systems) were used to detect change in level of cytokines in culture media. All reagents, kits and chemicals, unless otherwise stated, were used according to the manufacturers’ instructions. Other chemical unless specified were purchased from Sigma-Aldrich.

### Real-time quantitative PCR and NanoString profiling

RNA was extracted using RNeasy isolation kit (Qiagen). Real-time quantitative PCR (RT-qPCR) was then carried out using iTaq Universal SYBER Green One-Step kit (Bio-Rad) using primers specific for macrophage activation markers selected according to a previously published lists (23-25). Primers sequences are provided in (Supplemental Table S1) Results were normalized using 36B4 then expressed as fold change (FC) = 2^−ΔCt^, where ΔCt = (Ct_Target_ – Ct_36B4_) (24). For gene expression analysis by NanoString nCounter, cell lysates were hybridized to the 770 gene murine PanCancer Immune Profiling Panel according to the manufacturer’s protocol (NanoString Technologies). Briefly, 10 µl of Ambion Cells-to-Ct buffer (Thermo Fisher Scientific) was added to a cell pellet and a 5.0 µl volume of lysate was hybridized to the NanoString reporter and capture probes in a thermal cycler for 16 hr at 65°C. Washing and cartridge immobilization were performed on the NanoString nCounter PrepStation, and the cartridge was scanned at 555 fields of view on the nCounter Digital Analyzer. The resulting RCC files containing raw counts were reviewed for quality and normalized in the NanoString nSolver analysis software v3.0, followed by exportation and analysis.

### Flow cytometry and sorting protocol

Cells were collected, washed and incubated at 4°C in staining buffer (PBS, 2% BSA) containing the indicated surface antibodies. For intracellular staining, cells were fixed, permeabilized and stained using BD Cytofix/Cytoperm Fixation/Permeabilization kit (BD Biosciences) according to the manufacturer's instructions. Cells were then washed with staining buffer and subsequently analyzed. Data was recorded on a LSR II Flow Cytometer (BD Biosciences) and analysis completed using FlowJo software. Additional details are included in supplemental materials and methods.

### Western Blotting

Cell lysates with equal amounts of proteins (20-35 μg) were electrophoresed through 4–15% TGX Gel, then electrophoretically transferred to nitrocellulose membrane (Bio-Rad laboratories). Membranes were then incubated with the specified antibodies diluted according to the manufacturer's instructions. Membranes also were incubated with anti-α-tubulin or anti-GAPDH as a loading controls. Immunoreactive proteins were visualized with an appropriate peroxidase-conjugated secondary antibody.

### Confocal immunofluorescence

Macrophages cultured on chamber slides were washed twice with PBS, fixed in 3.8% formaldehyde for 20 min and permeabilized with 0.1% Triton X-100 for 5 min. Cells were washed twice with PBS, blocked with 2% BSA in PBS for 1 hr and subsequently incubated with CD206 antibody (1:800) at 4°C overnight. Cells were washed 3 times with PBS and incubated with appropriate fluorescent-labeled secondary antibodies at RT for 1 hr. Images were visualized using Leica TCS SP8 laser scanning microscope (Leica Microsystems).

### Histology and immunohistochemistry (IHC)

The histological specimens were embedded in paraffin, sectioned (4μm slices) and stained with haematoxylin & eosin (H&E). For immunohistochemistry, slides were stained using a Ventana Discovery XT automated system (Ventana Medical Systems). Briefly, slides were deparaffinized on the automated system with EZ Prep solution (Ventana). Enzymatic retrieval method was used in Protease 1 (Ventana). The rabbit primary antibodies that react to F4/80, α-SMA, CD206 (all purchased from Abcam) were used at a 1:400, 1:250, 1:1200 dilutions, respectively, in Dako antibody diluent (Agilent) and incubated for 60 min. The Ventana OmniMap Anti-Rabbit Secondary Antibody was used for 8 min. The detection system used was the Ventana ChromoMap kit, and slides were then counterstained with hematoxylin, followed by dehydrated and cover-slipping.

### Quantitative image analysis

Histology slides were scanned using the Aperio™ ScanScope XT with a 200X- (0.8NA) objective lens at a rate of 5 minutes per slide via Basler tri-linear-array. For TRAMP derived prostate tissue analysis, images and their meta-data were then imported into the Definiens Tissue Studio v4.0 suite. Each slide was then segmented into several tissue regions with stroma and gland being the main point of interest using the composer function in the software. The individual marker areas were then scored in terms of the intensity of F4/80, α-SMA and collagen. A pathologist was consulted to quality control that each tissue was correctly segmented into the regions of interest as shown in Supplemental Figure S4F. For CD206 frequency, images and their meta-data were imported into the Definiens Tissue Studio v4.2 suite. Slides were then analyzed by identifying individual cells using hematoxylin stain threshold and grown out to 2 μm. Cells were then identified by expression of IHC markers CD206 and F4/80. The segmented images were imported in Definiens Developer v2.4 and image contrast was used first to separate the tumor section from the background. Next, a 25 pixel ring was segmented around the periphery of the tumor to represent the edge of the tumor. Finally, the distance (in μm) to the nearest edge of tumor pixel was calculated for each cell in the image. Since each tissue section is a different size and shape, each distance to the edge value was normalized per mm^2^ of tissue. The normalized distances were then subject to histogram analysis to determine the percentage of cells that fall into 10 µm/mm^2^ bins representing areas of high macrophage abundance and higher acidity.

### TCGA PRAD analysis

The correlation of macrophage-related genes and glycolysis-related genes in a prostate cancer cohort was computed using level 3 gene expression estimates from the RNA-Sequencing in the TCGA PRAD (Prostate Adenocarcinoma Data Set) database, extracted, and hosted by Firehose DB (BROAD Institute, https://gdac.broadinstitute.org/). The expression estimates were derived using RSEM (Accurate transcript quantification from RNASeq) method (26). In Figures 1A and S1A, the original level 3 Illumina HiSeq RNAseqV2 RSEM gene-level normalized mRNA expression data for TCGA PRAD was downloaded from the TCGA data portal in March of 2016 and log2 transformed, log2(x+1). The 333 primary prostate tumors and associated clinical information, including reviewed Gleason score, were retrieved from the TCGA PRAD333 publication (27). Box and scatter plots were generated in MATLAB R2017a (MathWorks Inc.).

**Figure 1:**
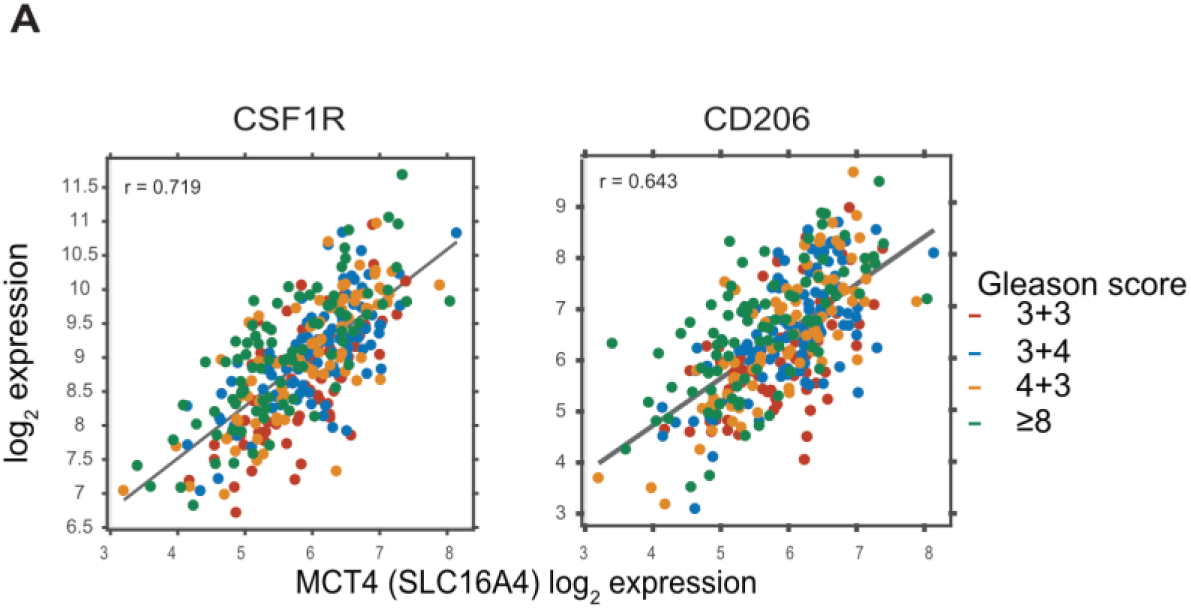
Macrophage infiltration correlates with MCT4 expression. **A)** Correlation between *CSF1R*/*CD206* and *MCT4* mRNA expression in early stage patients (Gleason score= 3+3) and advanced prostate (Gleason score = 3+4, 4+3 and ≥8) retrieved from TCGA PRAD333: *R*=0.719, 0.643, respectively.

### Agent Based Model

To examine the dynamics governing the interactions of macrophages and a metabolically aggressive tumor, we extended our previously published multiscale mathematical model that captures the complex spatiotemporal interactions of competing tumor cell phenotypes and microenvironmental selection forces such as oxygen, glucose, and acidosis (28, 29). Macrophages were added to this model to explore their ability to control tumor growth within this complex and dynamic environment. These macrophages can consume tumor and necrotic cells, as well as release and bind macrophage-derived cytokines. Macrophage behavior is modeled as a continuous phenotype from anti-tumor to pro-tumor like behaviors, determined by the local concentration of pH, pro- and anti-inflammatory cytokines, and the number of tumor and necrotic cells being digested (Figure 5A).

To calibrate the macrophage behavior, we used the statistical package R (30) to fit a linear model to specific gene expression data, in different ecological conditions, collected in the *in vitro* experiments described in Figure 2H. Each linear model takes the form

**Figure 2:**
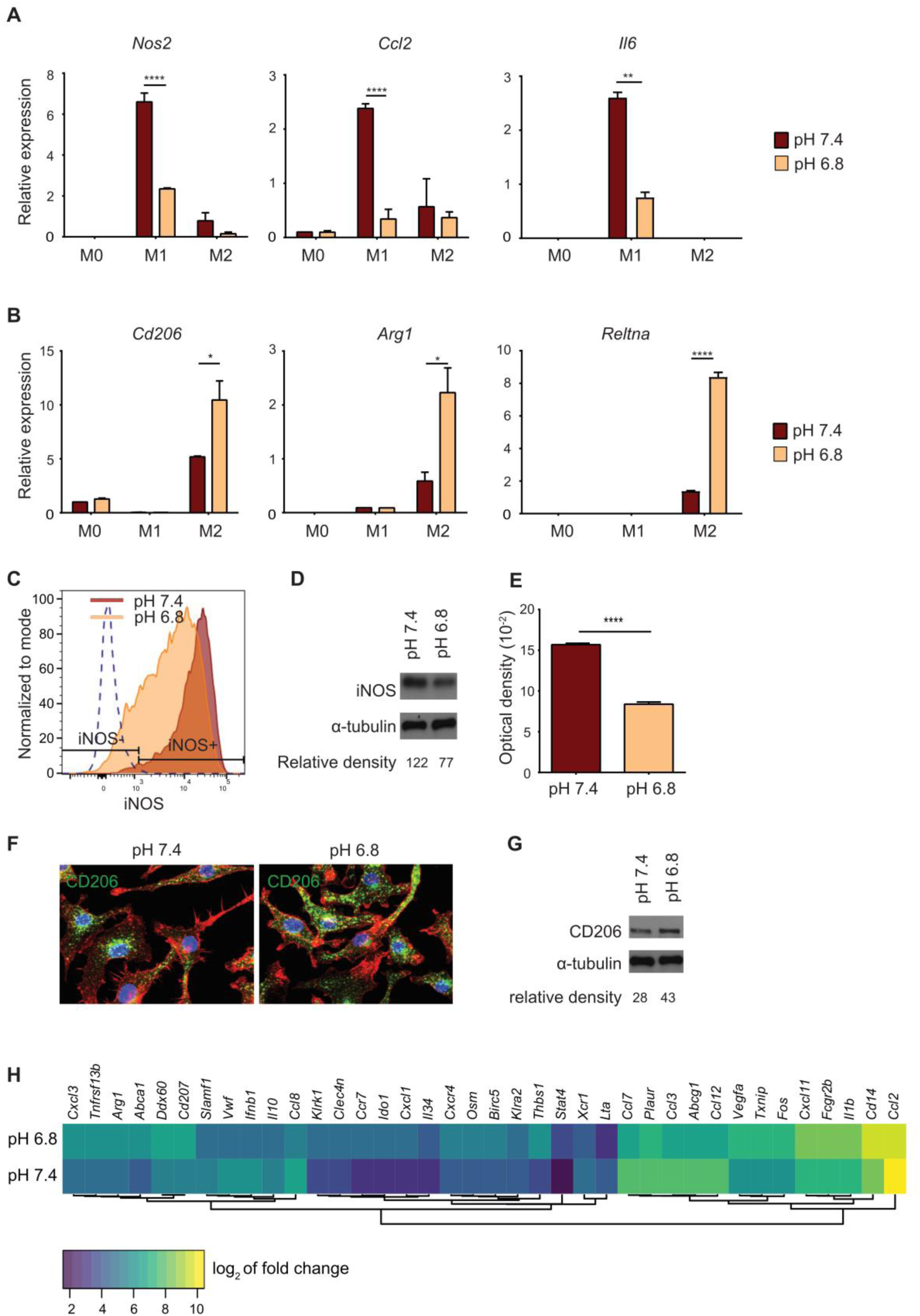
Extracellular acidosis alters macrophage activation *in vitro*. **A)** mRNA expression of *Nos2, Ccl2 and Il-6* **B)** mRNA expression of *Cd206, Arg1* and *Reltna* in BMDM stimulated for 24 hr with LPS/IFN-γ (M1) or IL-4 (M2), or left untreated (M0) at either pH 7.4 or pH 6.8. Data are presented as mean ± SEM. Two-way ANOVA was utilized for statistical analysis; *p<0.05, **p<0.01, ***p<0.001, ****p<0.0001. **C)** Expression of iNOS in LPS/IFN-γ activated macrophages at pH 7.4 or pH 6.8 using flow cytometry. **D)** Western blot analysis of iNOS in LPS/IFN-γ activated macrophages at pH 7.4 and pH 6.8. α-tubulin was used as a loading control. **E)** Nitrite level in supernatant of LPS/IFN-γ activated macrophages at pH 7.4 and pH 6.8, as measured by Griess reagent. Data are presented as mean ± SEM. Student’s t-test was utilized for statistical analysis; ****p<0.0001. **F)** Confocal immunofluorescent analysis of CD206 expression in IL-4 activated macrophages at pH 7.4 or pH 6.8. CD206 (green), Phalloidin (red) and Dapi (blue). **G)** Western blot analysis of CD206 expression in macrophages stimulated for 24 hr with IL-4 (M2) at pH 7.4 and 6.8. α-tubulin was used as loading control. **H)** Heatmap of the top differentially expressed genes (determined by *p* value and ranked by fold change) in LPS/IFN-γ activated macrophages at pH 7.4 or pH 6.8. nCounter PanCancer Immune Profiling that measure expression of 770 genes was used to assess difference in gene expression (n=2).

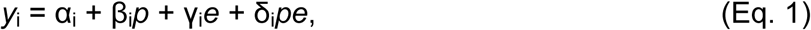

where y_i_ is the observed expression level of gene *i*, *p* is the pH, *e* represents the ecological conditions, and α_i_, β_i_, γ_i_, δ_i_ are determined during the fitting. In the model, the value of *e* for each cell at each time point is calculated using the equation

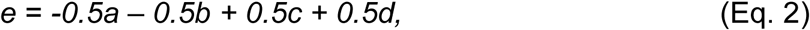

where *a* is the local inflammatory cytokine concentration, *b* is the number of tumor cells phagocytized, *c* is the local anti-inflammatory cytokine concentration, and *d* is the number of necrotic cells being phagocytized. During the fitting process, e is set to −1 for the inflammatory environment that promotes the extreme anti-tumor phenotype, while e is set 1 for the anti-inflammatory environment that induces the extreme pro-tumor phenotype. Thus, each macrophage checks the local extracellular pH, pro- and anti-inflammatory cytokine levels, number of tumor and necrotic cells being digested, and then adjusts each of the *i* phenotypic behaviors as dictated by the respective linear model. Further details are provided in the supplemental materials and methods.

### Quantification and statistical analysis

Unless otherwise indicated, unpaired two-tailed t-tests were used as appropriate for comparisons between two groups. Two-way ANOVA was used to compare multiple groups. Unless otherwise reported, GraphPad PRISM 7 software was used for statistical analysis. In TCGA data analysis, a two-sided Mann Whitney U test was used and median log_2_ fold change between the two groups was calculated. A significant change was defined when *p*<0.05 and log_2_ fold change > 0.585 (1.5x change). For the mathematical model, the Mantel-Haenszel test in the R package “survival” was used (31). To identify changes in macrophage phenotype using NanoString, differentially expressed genes with *p* <0.05 were ranked by fold-change with a cutoff of 1.5 or 2 (32). Statistical parameters including value of n, (mean ± SEM) and statistical significance and the tests used are reported in the figures and/or figure legends.

## Results

### Macrophage infiltration correlates with MCT4 expression

Advanced stages of prostate cancer adopt a high glycolytic phenotype that correlated with poor prognosis (33). The consequent lactic acid production was shown to aggravate highly immunosuppressive microenvironment through shaping macrophage phenotype in lung cancer and melanoma (15). Based on that, we questioned whether highly glycolytic phenotype correlates with macrophage infiltration or phenotype in late stage prostate cancer. Interestingly, analyzing publically available data of human prostate cancer revealed that *CSF1R* is expressed at higher levels in intermediate and late stage prostate cancers (Figure S1A). In addition*, CSF1R* and the macrophage activation marker *CD206* correlated with the monocarboxylate lactate transporter, *MCT4* (SLC16A4) in late stage prostate cancer (Gleason score 3+4, 4+3 and ≥8), as shown in Figure 1A and Supplemental Figures S1B, S1C. Of note, MCT4 facilitates lactate efflux and preserves intracellular pH by co-transporting lactate and protons across the plasma membrane of highly glycolytic and/or acid-resistant cells (16, 34). It is unknown whether the change in extracellular pH independent from changes in extracellular lactate concentration can modulate macrophage polarization in prostate cancer.

### Extracellular acidosis alters macrophage activation *in vitro*

Macrophages are highly plastic immune cells that display a range of phenotypic and functional properties (7, 35). To test whether an acidic tumor milieu can influence macrophage phenotype, we used zwitterionic buffer-based medium to stimulate bone marrow-derived macrophages (BMDM) using IFN-γ/LPS and IL-4, for 24 h at pH (7.4) or pH (6.8). Under these conditions, acidic pH did not affect viability of stimulated macrophages at 24 hr post-activation (Supplemental Figure S2A). As seen in Figures 2A and 2B, acidosis decreased gene expression of the pro-inflammatory markers *Nos2, Ccl2* and *Il-6* in IFN-γ/LPS -polarized macrophages, while it increased expression of anti-inflammatory markers *Cd206, Arg1* and *Reltna* in IL-4-polarized macrophages. Reduced iNOS protein levels were confirmed by flow cytometry and western blot (Figure 2C-2D). In line with the mRNA and protein expression data of iNOS, the level of nitrite in the culture media decreased, as shown in Figure 2E. Enhanced *Cd206* expression in IL-4-polarized BMDM was also confirmed by immunofluorescence and western blot (Figure 2F and 2G). Multi-analyte profiling in culture medium from these incubations also revealed significant alterations in the release of many inflammatory cytokines and chemokines (Supplemental Figures S2B and S2C). To expand these findings to other genes potentially involved in macrophage activation, we used NanoString profiling to assess the relative abundance of 770 cancer and immune related mRNAs. We observed that acidic pH increased the expression of a range of tumor associated macrophages (TAMs) related genes (e.g. *Arg1*, *Cd14, Il1b*) as well as angiogenesis associated genes (e.g. *Vegfa*, *Txnip*, *Thbs1*) in IFN-γ/LPS activated macrophages, in addition to a global decrease in the inflammation score (Figure 2H, Supplemental Figures S2D, S2E and Supplemental Table S2). These results demonstrate that extracellular acidosis alters macrophage activation towards a phenotype reminiscent of TAMs.

### Extracellular acidosis enhances a tumor-promoting macrophage phenotype

To examine if extracellular acidity could alter activation status of TAMs, we first activated BMDMs with tumor cell-conditioned medium at either pH 7.4 or 6.8. At pH 7.4, TRAMP-C2-conditioned medium significantly increased expression of *Arg1.* However, this effect was dramatically enhanced when the cells were activated at pH 6.8 (Figure 3A). Similarly, co-culturing BMDMs with TRAMP-C2 and TRAMP-C3 at pH 6.8 augmented *Cd206* mRNA expression and protein levels in macrophages as measured by RT-PCR and flow cytometry, respectively (Figures 3B, 3C, Supplemental Figures S3A, S3B). BMDM activated in acidic pH also increased the uptake of fluorescently-labeled ovalbumin, a mannosylated ligand endocytosed mainly through CD206 (Figure 3D). In addition, macrophage co-culture with TRAMP-C2 cells at acidic pH was associated with an increase in the release of inflammatory and angiogenic cytokines/chemokines (e.g. VEGF, CD14, M-CSF) known to be involved in tumor progression (Figure 3E).

**Figure 3:**
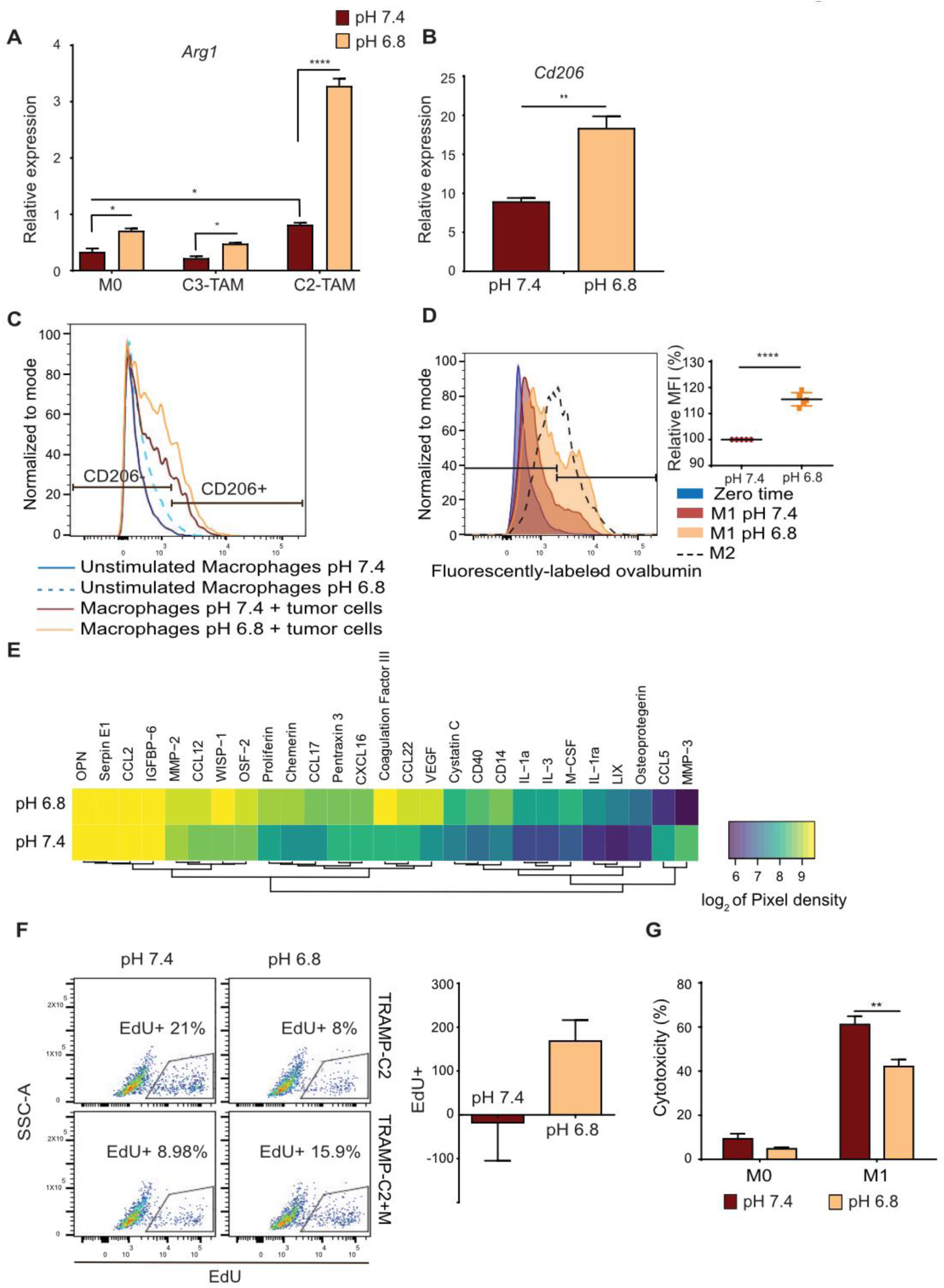
Extracellular acidosis enhances a tumor-promoting macrophage phenotype. **A)** Relative mRNA level of *Arg1* in macrophages treated with 30% TRAMP-C2, TRAMP-C3 conditioned medium at either pH 7.4 or pH 6.8 or left untreated as control (M0). Data are presented as mean ± SEM. Two-way ANOVA was utilized for statistical analysis; *p<0.05, ****p<0.0001. **B)** Relative mRNA level of *Cd206* in macrophages directly co-cultured with TRAMP-C2 for 4 days at pH 7.4 or pH 6.8, then sorted and processed for RNA extraction. Data are presented as mean ± SEM. Student’s t-test was utilized for statistical analysis; **p<0.01. **C)** Flow cytometry analysis of CD206 expression in macrophages incubated at pH 7.4 or pH 6.8 for 24 hr then either cultured alone or with TRAMP-C2 at pH 7.4 for another 24 hr. F4/80 staining was used to gate out tumor cells. **D)** Flow cytometry quantification of fluorescently-labeled ovalbumin uptake in LPS/IFN-γ activated macrophages at either pH 7.4 or 6.8 for 24 hr. Graph represents relative increase in fluorescently labelled ovalbumin uptake (n=5). Data are presented as mean ± SEM. Student’s t-test was utilized for statistical analysis; ****p<0.0001. **E)** Conditioned media from macrophage-tumor co-culture at pH 7.4 or pH 6.8 were processed for cytokines determination using mouse XL cytokine array. Densitometric analysis was then done using Image J software and pixel density was graphed as heatmap (n=2). **F)** TRAMP-C2 cells were co-cultured with or without macrophages in neutral or acidic medium for 24 hr. Cells were then labeled with EdU for 2 hr, collected and processed for flow cytometric analysis. SSC versus EdU fluorescence of TRAMP-C2 tumor cells in each culture condition was plotted. Fold change was calculated by dividing the EdU-incorporating cell count with macrophages by the corresponding values of tumor cells alone (n=6). **G)** Macrophages were activated with LPS/IFN-γ (M1) at pH 7.4 or 6.8 for 24 hr or left unstimulated as M0. Differentially activated macrophages were then co-cultured with TRAMP-C3 cells and lactate dehydrogenase in supernatants was measured 24 hr later to estimate cytotoxicity. Data are presented as mean ± SEM. Two-way ANOVA was utilized for statistical analysis; **p<0.01.

We next evaluated whether the phenotypic shift in macrophages would alter their function *in vitro.* TRAMP-C2 cells were incubated in acidic or neutral media in the presence or absence of non-polarized macrophages for 24 hr, and tumor cell proliferation was measured via EdU uptake after gating out F4/80^+^CD11b^+^ macrophages. As shown in Figure 3F, either acidic conditions or co-culture with unstimulated macrophages (pH 7.4) reduced tumor cell proliferation. In contrast, co-culturing with macrophages reversed the negative effect of acidic pH, resulting in a 2-fold increase in proliferation. The total number of cells was unchanged during the relative short period of the experiment (Supplemental Figure S3C). IFN-γ/LPS stimulated-macrophages are cytotoxic due to nitric oxide (NO) release, however, they lose their cytotoxic ability when if activated at low pH (Figure 3G). Acidic conditions therefore enhance a range of functions associated with the tumor-promoting phenotype of TAMs, at least *in vitro*.

### Buffering tumor-secreted acids alters TAM phenotype *in vivo* and reduces tumor progression

To determine whether tumor acidity was a contributing factor to the phenotype of TAMs, we first used mice subcutaneously implanted with TRAMP-C2 cells, which grow fast in the subcutaneous setting (4 weeks) and thus are refractory to the chemopreventive effect of systemic buffering reported earlier in transgenic models (Supplemental Figures S4A – S4C). This provided us the opportunity to evaluate whether tumor acidity had a direct impact macrophage phenotype under constant tumor volume. In addition, when we treated tumor bearing mice with oral buffer (200 mM ad lib NaHCO_3_) to neutralize tumor acidity, analysis of myeloid cell infiltration by flow cytometry revealed no significant differences (Supplemental Figure S4D). This provided another opportunity to test polarizing effect of acidity independent from changes in the number of immune cells. Accordingly, we then analyzed the impact of buffering tumor acidity on macrophage activation using NanoString profiling and RT-PCR quantification of selected genes in sorted TAMs. As shown in Figure 4A, buffering tumor acidity increased the NanoString-derived “inflammation score”, denoting a shift towards a anti-inflammatory phenotype. There were also decreases in the expression of major TAMs markers including *Arg1* and *Fcgr2b* (Figure 4B, Supplemental Table S3). In a separate set of experiments, we also observed a significant reduction in *Cd206* and *Arg1* by single reaction RT-PCR (Figure 4C). In agreement with this, quantitative image analysis of formalin-fixed sections showed a significant drop in the density of CD206 positivity in bicarbonate-treated tumors compared to untreated controls (Supplemental Figure S4E). We second examined the TRAMP transgenic prostate model, which allowed us to test the effect of buffering tumor acidity over extended time scale (32 weeks). In this model, macrophage infiltration but not SMA^+^ fibroblasts corresponded with tumor progression, with the highest infiltration coincident with loss of fibromuscular tunica, disease progression from PIN lesions to high-grade adenocarcinomas, and invasion (Figures 4D-4F). Algorithm generated segmentation used to quantify those cell types is shown in Supplemental Figure S4F. In addition, representative images are shown in Supplemental Figure S4G. To investigate the role of pH, we treated TRAMP mice with 200 mM ad lib NaHCO_3_ for 28 weeks, starting at 4 weeks of age. Prostate tissue isolated from buffered TRAMP mice showed lower infiltration of F4/80^+^ macrophages into the stromal compartment compared to controls (Figure 4G and S4H). Furthermore, buffering tumor pH normalized prostate interglandular structure, decreased the relative percentage of the stromal compartment, and reduced tumor incidence as compared with control (Figure 4H and 4I). Together these results indicate that the acidic microenvironment contributes to the pro-tumor polarization state of TAMs as well as tumor progression.

**Figure 4:**
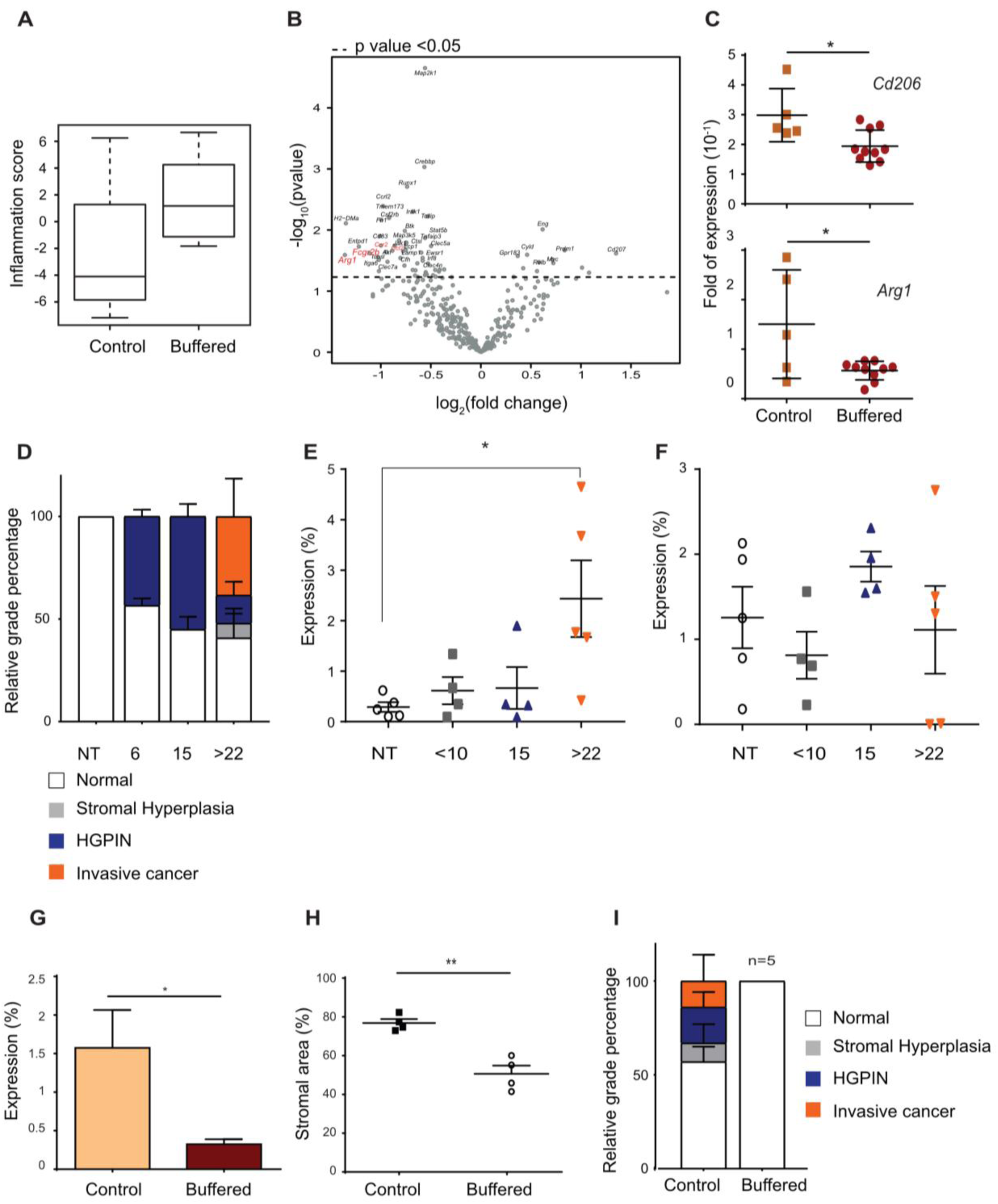
Buffering tumor-secreted acids alters TAM phenotype *in vivo* and reduces tumor progression. **A)** Inflammation score and **B)** Volcano plot generated by nSolver software 3.0 using gene expression data from nCounter PanCancer Immune profiling of CD11b^+^F4/80^+^ tumor associated macrophages (TAMs) sorted from control or sodium bicarbonate-treated (buffered) TRAMP-C2-bearing mice (n=4-5). **C)** Fold expression of *Cd206* and *Arg1* in CD11b^+^F4/80^+^ TAMs sorted from independent cohort of control or sodium bicarbonate-treated (buffered) TRAMP-C2-bearing mice (n=5-10). **D)** Histopathological analysis of H&E samples from 6, 15, 22, 23 and 25 weeks old TRAMP mice. **E, F)** Quantification of F4/80 (macrophage) and α-SMA (fibroblast) staining in serial sections of paraffin embedded prostates, isolated from 6, 15, 22, 23 and 25 weeks old TRAMP mice. **G)** TRAMP mice were treated with sodium bicarbonate buffer starting from 4 weeks of age (buffered) or kept on tap water as control. F4/80 stained sections were digitally quantified and percentage of F4/80 staining intensity in stroma of prostate tissue were plotted (n>4). **H)** Mean area of segmented stromal compartment (n=4). **I)** Histopathological analysis of H&E slides of buffered and control TRAMP mice. Data are presented as mean ± SEM. Student’s t-test was utilized for statistical analysis; *p<0.05, **p<0.01.

### Acid-responsive macrophages promote tumor growth *in silico*

Despite the impact of buffering on prostate carcinogenesis and its impact on the phenotype of TAMs, it was unclear whether these were functionally related, as acidic pH is thought to impact a range of biological processes within tumors. To test whether acid-responsive macrophage can enhance tumor progression, we developed an *in silico* agent based model (Figure 5A) that can turn off macrophage acid-induced responses regardless of the underlying mechanisms In this model, tumor acidity emerges from increased glycolytic metabolism in combination with poor perfusion, and it affects macrophage phenotype as modulated between two extremes states (Figure 5B). Two scenarios were imposed in order to determine the impact of pH on the ability of a constant number of macrophages to modulate tumor growth. In the first scenario, macrophages behave phenotypically as if they are in pH 7.4 regardless of the actual local pH value (i.e. the value of *p* in Eq. 1 is set to 7.4 regardless of the actual local pH). In the second scenario, macrophage behavior is modulated by setting *p* in Eq. 1 to the local pH calculated at that position in the model. Simulations were run until either 1) the tumor took over 90% of the domain, or 2) ten simulated years had elapsed, indicating that the tumor had successfully been eradicated or controlled. Each scenario was run 100 times, and the time of 90% takeover was recorded at the end of each run. As shown in representative simulation images (Figure 5C), the extracellular acidosis, created by excess tumor glycolysis, dynamically changes the macrophage phenotype represented by *Arg1* and *Ccl2* expression. The time to tumor takeover can be visualized using Kaplan-Meier curves (Figure 5D). The tumors grew much more rapidly in the simulations where acidosis was actively modulating macrophage behavior. The difference in these survival curves was significant, with p=1.1×10^-9^ as calculated using the Mantel-Haenszel test. The results from these simulations suggest that acid released by tumor cells can create a protective niche capable of directing the functional role of macrophages, thereby increasing tumor growth and decreasing time to progression.

**Figure 5:**
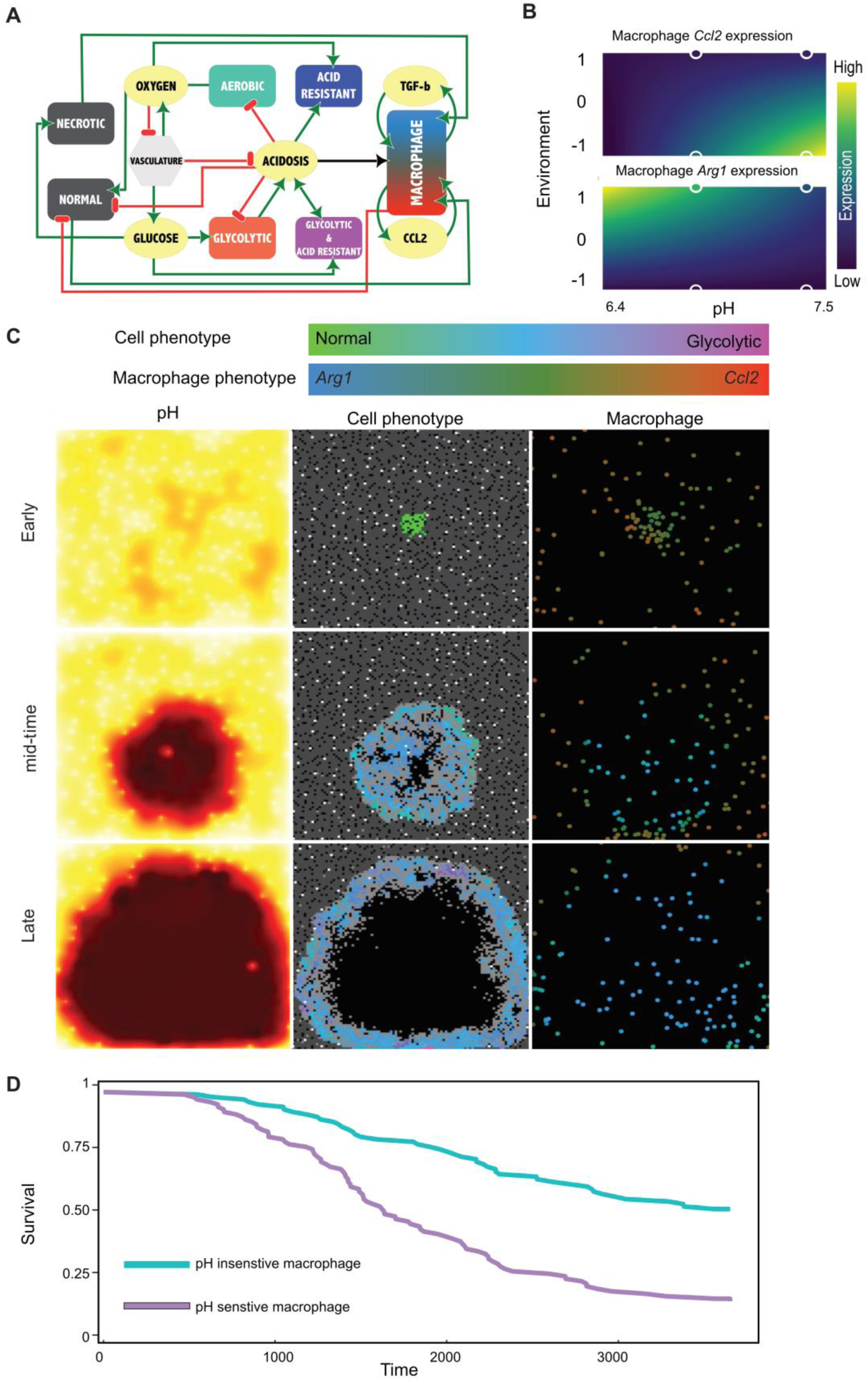
Acid-responsive macrophages promote tumor growth *in silico*. **A)** Interaction network for agent based model illustrating how macrophages and cells interact with, and are affected by, the microenvironment, which is composed of glucose, oxygen, acid, necrotic cells, and pro- and anti-inflammatory cytokines. Green lines reflect promotion, while red lines indicate inhibitory interactions. **B)** Output of linear model fitting of *Arg1* and *Ccl2* expression represented as heatmap. For each phenotypic trait, a linear model allows to predict expression under a variety of conditions. Here, −1 is a tumor rich inflammatory environment, while 1 is environment with necrosis and anti-inflammatory cytokines. The circles outlined in white are the actual *in vitro* data. **C)** Snapshots from agent based model. In the pH window, low pH is dark red, while high pH is yellow. In the cell window, grey pixels are normal cells, white are vessels, and tumor cells are colored by their phenotype. In the macrophage window, each macrophage is colored by the mix of of CCL2 and ARG1 expressed. Top panel is early in the simulation, bottom panel is when the tumor has taken over 90% of the domain and the simulation is stopped. **D)** Simulated survival curves generated after running the simulation under two scenarios, one hundred times each. The “pH Insensitive Macrophage” scenario is where macrophages are not affected by pH, while macrophage behavior is affected by acid in the “pH Sensitive Macrophage” scenario. Here, survival time is the amount of time it took the tumor to take over 90% of the domain, given a maximum amount of time of 10 years. The Mantel-Haenszel reveals that these survival curves are significantly different, with p=1.1e-9.

## Discussion

Tumors undergo metabolic transformation that rewires cellular metabolism to promote tumorigenicity, immune evasion and disease recurrence (36). One of these metabolic abnormalities is upregulation of glycolysis, even under aerobic conditions. High rate of glycolysis provides malignant cells with proliferative privilege by facilitating uptake and incorporation of nutrients into the growing biomass (37). Metabolic byproducts of glycolysis, such as lactic acid, also cause a heterogeneous acidification of the extracellular space, which can results in immunosuppressive nature of the tumor microenvironment (^6^, ^38)^. Unlike studies that combine lactate and H+ ions as single functional entity named “lactic acid”, we identified an independent role of tumor-generated acidity in driving TAMs phenotype, which in turn can contribute to tumor progression.

In the current investigation, we propose a scenario in which acids generated by glycolytic cells alter the phenotype of TAMs, creating a permissive niche for cancer progression in prostate cancer. Using zwitterionic organic chemical buffering system, our data shows that acidic pH alter the activation state of macrophages incubated under polarizing conditions, directing the cells towards a functional state similar to the pro-tumor phenotype often ascribed to TAMs. Furthermore, we demonstrate that buffering tumor acidosis alters the activation state of TAMs, with a significant reduction in genes such as *Arg1* and *Cd206* that are usually associated with a tumor-promoting role for this population. Finally, we noted an association between tumor progression, acidosis, and the presence of macrophages in prostate cancer progression in mice and human disease, and utilize an *in silico* agent-based model to delineate a role for acidosis in regulating macrophage phenotype and tumor progression. Cumulatively these results suggest that tumor acidosis is an important factor that dictates the pro-tumor functionality of macrophages in prostate cancer.

Lactic acid produced by tumor cells was reported earlier to polarize macrophages into an M2-like phenotype, with *Arg1* expression by macrophages essential for lung cancer and melanoma growth(15). In addition, Carmona-Fontaine *et al*. have demonstrated that lactate cooperates with hypoxia to induce the expression of ARG1 in macrophages. Through the employment of an agent-based model, they also showed that hypoxia-responsive macrophages induce faster tumor growth (39). However, there is limited information regarding how acidity, independent from those metabolic factors, can influence properties of macrophages in tumor microenvironment. Only recently, Toszka *et al*. identified a role of tumor acidity independent from lactate in driving growth of melanoma cell line B16 in cAMP dependent manner (40), In the current study that investigates prostate cancer, we also provide evidence that acidic pH, independent from lactate can promote the pro-tumor polarization of macrophages, including enhanced tumor cell proliferation, loss of cytotoxicity, and release of angiogenic factors. *In silico* modeling also demonstrated that modulation of the macrophage phenotype by acidity was a significant driver of tumor progression. Overall, our findings suggest that tumor progression could be reduced by intervening with the acid-induced phenotypic changes of macrophages in the tumor microenvironment.

## Supporting information

## Acknowledgments

We thank Agnieszka Kasprzak and Dominique Abrahams for technical assistance. We also thank Bailey Philips from Moffitt’s summer high school internship program in integrated mathematical oncology (HIP-IMO), during which part of the *in silico* model was developed. This work was supported by the Moffitt Cancer Center Flow Cytometry, Molecular Genomics, Analytic Microscopy, and Tissue Core Facilities, all comprehensive cancer center facilities designated by the National Cancer Institute (P30CA076292).

## Contributions

A.E. and R.G. conceived the idea and designed the study. A.E. designed and conducted experiments. A.E., B.R. and R.G analyzed and interpreted the data. C.G., M.R. and R.B. designed, built, tested and wrote the mathematical section of the manuscript. Y.B. and A.B. performed the analysis of publically available TCGA data. J.D. performed the IHC assessment. K.L., N.V., and A.I. assisted with experiments and/or optimizing protocols. A.E. wrote and compiled the manuscript in consultation with R.G. and B.R. All authors contributed to the manuscript. J.C., R.G., S.P, A.A., B.R. and R.G. supervised the project.

## Data and software availability

The authors declare that the main data supporting the findings of this study are available within the article and its Supplementary Information files.

