## Supplementary material for "Acidity promotes tumor progression by altering macrophage phenotype in prostate cancer"

**Supplemental tables**

**Table S1**

**Primers sequences of major M1 and M2 markers of mouse macrophages**

| ***36B4*** | **GCTCCAAGCAGATGCAGCA**  **CCGGATGTGAGGCAGCAG** |
| --- | --- |
| ***β-actin*** | **CTAAGGCCAACCGTGAAAAG**  **GGTACGACCAGAGGCATACA** |
| ***Cd206 (Mrc1)*** | **CAGGTGTGGGCTCAGGTAGT**  **TGTGGTGAGCTGAAAGGTGA** |
| ***Arg1*** | **TTTTTCCAGCAGACCAGCTT CATGAGCTCCAAGCCAAAGT** |
| ***Mcp1*** | **GGGATCATCTTGCTGGTGAA**  **AGGTCCCTGTCATGCTTCTG** |
| ***iNos*** | **TCCAGGGATTCTGGAACATT**  **GAAGAAAACCCCTTGTGCTG** |
| ***il-6*** | **AACGATGATGCACTTGCAGA** **TGGTACTCCAGAAGACCAGAGG** |
| ***Retnla*** | **GGGATAGTTAGCTGGATTGGC**  **CCTTCTCATCTGCATCTCCC** |

**Table S2**

**Differentially expressed mRNA of M1 macrophages activated at pH 6.8 compared with pH 7.4 (filtered by fold change and p value)**

|  | **mRNA** | **Log_2_ fold change** | ***p* value** |
| --- | --- | --- | --- |
| 1 | *Thbs1* | 3.35 | 0.000902 |
| 2 | *Abca1* | 2.41 | 0.00263 |
| 3 | *Cxcl1* | 1.95 | 0.0216 |
| 4 | *Il1b* | 1.88 | 0.0085 |
| 5 | *Arg1* | 1.85 | 0.00553 |
| 6 | *Cd14* | 1.76 | 0.000505 |
| 7 | *Ccr7* | 1.75 | 0.0169 |
| 8 | *Birc5* | 1.71 | 0.0329 |
| 9 | *Vegfa* | 1.68 | 0.00269 |
| 10 | *Stat4* | 1.67 | 0.00248 |
| 11 | *Fos* | 1.64 | 0.00337 |
| 12 | *Tnfrsf13b* | 1.56 | 0.0346 |
| 13 | *Ido1* | 1.54 | 0.0172 |
| 14 | *Fcgr2b* | 1.51 | 0.00228 |
| 15 | *Cxcr4* | 1.43 | 0.00786 |
| 16 | *Clec4n* | 1.36 | 0.00168 |
| 17 | *Il34* | 1.32 | 0.00293 |
| 18 | *Cxcl11* | 1.31 | 0.00675 |
| 19 | *Cd207* | 1.25 | 0.00614 |
| 20 | *Ddx60* | 1.25 | 0.00637 |
| 21 | *Cxcl3* | 1.24 | 0.00691 |
| 22 | *Txnip* | 1.2 | 0.00174 |
| 23 | *Osm* | 1.14 | 0.0234 |
| 24 | *Klrk1* | 1.06 | 0.0138 |
| 25 | *Klra2* | 1.04 | 0.041 |
| 26 | *Il10* | -1.15 | 0.00147 |
| 27 | *Plaur* | -1.16 | 0.00944 |
| 28 | *Slamf1* | -1.17 | 0.0158 |
| 29 | *Ccl2* | -1.29 | 0.000135 |
| 30 | *Ccl7* | -1.32 | 0.000511 |
| 31 | *Xcr1* | -1.33 | 0.000583 |
| 32 | *Ifnb1* | -1.39 | 0.00905 |
| 33 | *Vwf* | -1.42 | 0.00326 |
| 34 | *Lta* | -1.43 | 0.0121 |
| 35 | *Syt17* | -1.44 | 0.0449 |
| 36 | *Ccl3* | -1.52 | 0.00258 |
| 37 | *Ccl8* | -1.64 | 0.0126 |
| 38 | *Abcg1* | -1.74 | 0.000949 |
| 39 | *Ccl12* | -1.79 | 0.0015 |
| 40 | *Tnfsf14* | -2.08 | 0.00921 |

**Table S3**

**Differentially expressed mRNA of TAMs isolated from buffered compared with TAMs isolated from control mice (filtered by both fold change and *p* value)**

|  | **mRNA** | **Log_2_ fold change** | ***p* value** |
| --- | --- | --- | --- |
| 1 | *Arg1* | -1.36 | 0.0258 |
| 2 | *H2-DMa* | -1.35 | 0.00782 |
| 3 | *Entpd1* | -1.22 | 0.0188 |
| 4 | *Fcgr2b* | -1.11 | 0.0243 |
| 5 | *Icosl* | -1.03 | 0.0283 |
| 6 | *Itga6* | -1.02 | 0.031 |
| 7 | *Tnfrsf14* | -1.02 | 0.0474 |
| 8 | *Tnfrsf14* | -1.02 | 0.0474 |
| 9 | *Cd83* | -1.01 | 0.0126 |
| 10 | *Ccr2* | -1.00 | 0.0181 |
| 11 | *Fn1* | -0.996 | 0.00688 |
| 12 | *Ccrl2* | -0.979 | 0.00408 |
| 13 | *Axl* | -0.944 | 0.023 |
| 14 | *Clec7a* | -0.934 | 0.0334 |
| 15 | *Tmem173* | -0.919 | 0.0064 |
| 16 | *Csf2rb* | -0.916 | 0.0062 |
| 17 | *Ccl2* | -0.886 | 0.022 |
| 18 | *Jak1* | -0.863 | 0.0183 |
| 19 | *Map3k5* | -0.819 | 0.0149 |
| 20 | *Cfh* | -0.805 | 0.029 |
| 21 | *Lamp1* | -0.777 | 0.0225 |
| 22 | *Irf5* | -0.765 | 0.0385 |
| 23 | *Btk* | -0.761 | 0.0104 |
| 24 | *Lcp1* | -0.752 | 0.0212 |
| 25 | *Ctsl* | -0.749 | 0.0174 |
| 26 | *Tnfaip3* | -0.745 | 0.0162 |
| 27 | *Runx1* | -0.738 | 0.00197 |
| 28 | *Ifngr1* | -0.682 | 0.0454 |
| 29 | *Irak1* | -0.681 | 0.005 |
| 30 | *Vhl* | -0.654 | 0.0461 |
| 31 | *Itch* | -0.636 | 0.0354 |
| 32 | *Ewsr1* | -0.596 | 0.0236 |
| 33 | *Irf8* | -0.583 | 0.0289 |
| 34 | *Clec4n* | -0.582 | 0.032 |
| 35 | *Crebbp* | -0.565 | 0.00094 |
| 36 | *Map2k1* | -0.559 | 2.25E-05 |
| 37 | *Stat5b* | -0.559 | 0.0136 |
| 38 | *Cd48* | -0.552 | 0.0494 |
| 39 | *Cd48* | -0.552 | 0.0494 |
| 40 | *Tollip* | -0.533 | 0.00602 |
| 41 | *Havcr2* | -0.529 | 0.0356 |
| 42 | *Clec5a* | -0.499 | 0.0184 |
| 43 | *Psma2* | -0.457 | 0.0454 |
| 44 | *Cd63* | -0.42 | 0.0482 |
| 45 | *Cd63* | -0.42 | 0.0482 |
| 46 | *Tgfb1* | -0.412 | 0.045 |
| 47 | *Eng* | 0.616 | 0.00983 |
| 48 | *Myc* | 0.713 | 0.0306 |
| 49 | *Tnfrsf11a* | 0.723 | 0.035 |
| 50 | *Prdm1* | 0.836 | 0.0216 |
| 51 | *Vim* | 1.01 | 0.0416 |
| 52 | *Cd207* | 1.35 | 0.0243 |

**Table S4**

**List of genes curated in the glycolytic pathway (HSA00010, KEGG:** [**http://www.genome.jp/dbget-bin/www_bget?hsa00010**](http://www.genome.jp/dbget-bin/www_bget?hsa00010)**)**

| **KEGG Pathway (Glycolysis)** | | | | |
| --- | --- | --- | --- | --- |
| ACSS1 | ALDH3A1 | ENO2 | LDHA | PGAM1 |
| ACSS2 | ALDH3A2 | ENO3 | LDHAL6A | PGAM2 |
| ADH1A | ALDH3B1 | FBP1 | LDHAL6B | PGAM4 |
| ADH1B | ALDH3B2 | FBP2 | LDHB | PGK1 |
| ADH1C | ALDH7A1 | G6PC | LDHC | PGK2 |
| ADH4 | ALDH9A1 | G6PC2 | PCK1 | PGM1 |
| ADH5 | ALDOA | GALM | PCK2 | PGM2 |
| ADH6 | ALDOB | GAPDH | PDHA1 | PKLR |
| ADH7 | ALDOC | GCK | PDHA2 | PKM2 |
| AKR1A1 | BPGM | GPI | PDHB | TPI1 |
| ALDH1A3 | DLAT | HK1 | PFKL |  |
| ALDH1B1 | DLD | HK2 | PFKM |  |
| ALDH2 | ENO1 | HK3 | PFKP |  |
