## Supplementary material for "Acidity promotes tumor progression by altering macrophage phenotype in prostate cancer"

**Supplemental Materials and Methods**

**Flow cytometry and sorting protocol**

Tumor cell suspensions were prepared from solid tumors using digestion buffer (0.5 mg/ml collagenase I, 0.5 mg/ml collagenase IV, 0.25 mg/ml hyaluronidase V, 0.1 mg/ ml DNase I in Hanks balanced salt solution (HBSS)) with constant stirring for 2 hours at room temperature. The resulting suspension was passed through a 70-µm cell strainer and washed once with HBSS. Cells were resuspended in staining buffer (PBS, 2% BSA) to a concentration of 0.5−1X10^6^ cells/ml for flow cytometric analysis. For sorting experiments, 1-5X10^7^ cells were slowly cryopreserved in freezing medium (10% DMSO and 90% (DMEM, 1% BSA)) and stored in -80 ºC to be sorted later on single day. Staining protocol involves incubation with live/dead (L/D) fixable fluorescent reactive dyes (Molecular Probes) to exclude dead cells. After washing, cells were incubated for 30 min in 100 μl staining buffer containing the following antibodies against surface markers: CD45, CD11b, Ly6C, F4/80, MHCII, Ly6G and/or CD206. Cells were then washed and subsequently sorted or analyzed by flow cytometry. A representative example of sorting gating strategy is shown in supplemental Figure S6A. For experiments that involve intracellular markers, cells were fixed, permeabilized and stained using BD Cytofix/Cytoperm Fixation/Permeabilization kit (BD Biosciences) following manufacturer's instructions.

For ovalbumin uptake, activated macrophages were starved for 2 hr using FBS-free medium, then treated on ice with 20 µg/ml DQ ovalbumin (Molecular Probes) in ice-cold neutral FBS-free medium. Treated macrophages were rapidly transferred to incubator for 25-50 min at 37 ºC or left at 4ºC for Zero time measurement. Macrophages were then washed twice with ice cold FBS-free medium, washed once with PBS, collected using Cellstripper non-enzymatic cell dissociation solution (Corning), rapidly fixed using 4% formaldehyde in PBS, washed then analyzed by flow cytometry.

For EdU proliferation assay, tumor cells were co-cultured with macrophages for 24 hr then EdU substrate was added to culture medium in a final concentration of 10 µM for another 2 hr at 37 ºC. Macrophages were then washed three times with PBS, collected using Cellstripper non-enzymatic cell dissociation solution (Corning). Cells were then fixed, permealized and stained for incorporated EdU following manufacturer’s instructions. PE-conjugated F4/80 and PE-Cy7-conjugated CD11b staining were then performed to exclude macrophage from analysis.

**Additional details for the mathematical model:**

To examine the dynamics governing the interactions of macrophages and metabolically aggressive tumors, we have extended our previously published multiscale mathematical model (1, 2). Briefly we outline here the previously published model, and following we describe the addition of macrophages specific to this work.

The mathematical model is a two-dimensional hybrid cellular automaton (HCA) that can capture the complex spatiotemporal interactions of competing tumor cell phenotypes and microenvironmental selection forces such as oxygen, glucose, and acidosis. The core elements are shown in Figure 5A, and the parameter values can be found in Table S5.

Diffusion of oxygen, glucose and protons is modeled with a set of reaction-diffusion partial differential equations (PDEs) that is solved using an ADI scheme (3) for speed. Due to the interior boundary conditions imposed by the point-source vasculature, we used a low Courant number of 0.2 for stability. Typical convergence to a relative tolerance of 10^-4^ was achieved in less than 100 steps of the ADI, typically 20-30 steps once the transients from initial conditions dissipated. The domain boundary conditions for oxygen, glucose, and protons were set to the average concentration of the diffusibles in normal tissue, in the absence of a tumor. Specifically, a normal tissue with vasculature was initialized and advanced through the simulation, updating the boundary condition to match the average value of the diffusible concentrations across the entire domain. Once all the cells and concentrations reached relaxed into dynamic equilibrium, those values then served as the fixed boundary concentrations for the simulations performed with a tumor.

The agent based model (ABM) was simulated on a square grid with dimensions of 100 by 100. The algorithm loops through the cell positions in a random order to avoid bias when determining cell behaviors such as proliferation. Cells can only proliferate into empty space if they have completed a cell cycle. The cell cycle is advanced by a sigmoidal function of the ATP production rate (f_A_), with higher ATP rates leading to faster progression through the cycle (up to a limit). Once the cycle is complete, the cell can proliferate on any subsequent step of the ABM, when space is available. If f_A_ is less that the quiescent threshold A_q_, the cycle is not advanced on that step. If f_A_ is less than the death threshold A_d_, then the cell has a chance of becoming necrotic on that step. Necrotic cells occupy space and degrade into empty space with rate p_n_ but do not have any metabolic function.

Initially, blood vessels are seeded into the tissue using a circle packing algorithm, with an average density taken from histological sections. Vessels are not placed closer than the minimum distance of σ_min_. Angiogenesis is simulated by probabilistically adding vessels to hypoxic areas based on the size of the area, with larger areas increasing the probability that a new vessel will be placed randomly within the hypoxic zone. Naturally, vessels added as such will begin to deliver oxygen and lessen the hypoxia; this serves to slow the angiogenic process as normal vascular density is reached. Vessels are degraded by growing tumor cells as follows. New vessels are seeded with a vessel stability parameter v_mean_. If a tumor has no space to divide on a time step, and it is next to a vessel, it will lower the value of this parameter by 1. If that parameter for that vessel reaches 0, it is removed from the simulation. By adjusting the strengths of the angiogenic probabilities and the vessel stability parameter, tumors with different vascular dynamics can be simulated.

In silico, macrophage behavior is based on the patterns of gene expression observed in the *in vitro* experiments (Results section 1). *In vitro* macrophages polarized to an M1 state are assumed to represent the extreme end of the M1 phenotype. In the model, this would be the equivalent of being saturated in inflammatory cytokines and having phagocytized the maximum number of tumor cells by setting a= 1, b=1, c=0, d=0, yielding e=-0.5a -0.5b + 0.5c + 0.5d = ‑1. A similar assumption is made about experimentally polarized M2 macrophages, that is, they represent the extreme end of M2 expression levels. This is modeled by setting a=0, b=0, c=1, d=1, yielding e=-0.5a -0.5b + 0.5c + 0.5d = 1.

Given the above assumptions, a linear model can be fit to the expression levels of any gene of interest in the four experimental conditions (see Table S6). Given each linear model, it is then possible to predict what expression would be under a continuous range of conditions (Supplemental Figures S5A-S5G). Here, the range of pH values modeled is 6.3-7.5.

In the model, a, b, c, and d are all scaled between 0 and 1 before calculating e. The minimum and maximum values used in the scaling can be found in (Table S7), and are obtained from the fits described above. These minimum and maximum values occur when either pH is 6.3 or 7.5. During the simulation, any scaled value below 0 is set to 0, while scaled values greater than 1 are set to 1.

Diffusion of macrophage-derived cytokine is also modeled using the ADI scheme, as described above. A difference here is that cytokines decay over time and are absorbed by the vasculature. Decay is modeled by multiplying the cytokine diffusion field by the decay rate, λ, during each iteration of the diffusion loop. Vascular absorption is simulated by setting the concentration of cytokine to zero at each vessel.

The binding of cytokine to cell receptors is modeled using the Hill equation with the Hill coefficient being 1 (supplemental Figure S5H). Following diffusion, macrophages detect the local concentration of cytokine and, using a disassociation constant, determine the fraction of their receptors that have bound the cytokine. The actual amount of cytokine bound is found by multiplying this value by the number of receptors the macrophages currently expresses. This amount of cytokine is then used to determine a, in the case of CCL2, and c, in the case of TGF-β.

Macrophage movement is modeled as a combination of random movement and chemotaxis in response to the gradient of the CCL2 diffusible. If we define the diffusible concentration at the lattice position that the macrophage currently occupies as C(x,y), then the X component of the gradient is computed as C(x+1,y)-C(x-1,y) and the Y component of the gradient is computed as C(x,y+1)-C(x,y-1). If some of these components fall outside the grid domain, then these values are substituted with C(x,y). The norm of this gradient is the strength of the effect of chemotaxis on the movement of the macrophage. We modulate this norm by dividing by the strength of chemotaxis, θ. If the norm is zero, then the cell just moves to a random position chosen uniformly from a circle of radius ψ around the macrophage. If the norm is nonzero, then we construct the movement circle at a position norm distance away from the macrophage, with displacement along the gradient. We then sample this circle to get the next movement position in the same fashion as when the norm is zero. This simulates the combination of chemotaxis and random movement.

**Table S5**

**Mathematical model parameters**

| **Parameter** | **Value** | **Units** | **Description** | **Reference** |
| --- | --- | --- | --- | --- |
| δx | 20 | μm | Diameter of CA grid point  dsd | Model specific |
| p_D_ | 0.005 | 1/day | Normal cell death rate | (4) |
| p_δ_ | 0.7 | 1/day | Prob. of death in poor conditions | Model specific |
| p_n_ | 5x10^-4^ | 1/day | Necrotic turnover rate | Model specific |
| D_O_ | 1820 | μm^2^/s  μm^2^/s | Diffusion rate O_2_ | (5) |
| D_G_ | 500 | μm^2^/s | Diffusion rate G | (6) |
| D_H_ | 1080 | μm^2^/s | Diffusion rate protons | (7) |
| O_o_ | 0.056 | mM | O_2_ concentration in vessels | (8) |
| G_o_ | 5 | mM | G concentration in vessels | (9) |
| pH_o_ | 7.4 | pH | Blood vessel pH | Model specific |
| V_O_ | 0.012 | mM/s | Max O_2_ consumption | (8) |
| k_O_ | 0.005 | mM | Half-max O_2_ consumption | (10) |
| k_G_ | 0.04 | mM | Half-max G consumption | (10) |
| k_H_ | 2.5x10^-4^ | – | Buffering parameter | (7) |
| β_N_ | 6.65 | pH | Normal cell acid resistance | (11) |
| β_T,min_ | 6.1 | pH | Max tumor cell acid resistance | (10) |
| p_G,max_ | 50 | – | Max glycolytic parameter | (12) |
| ∆H | 0.003 | – | Drift rate acid resistance | Model specific |
| ∆G | 0.15 | – | Drift rate glycolysis | Model specific |
| A_d_ | 0.3 | – | ATP rate threshold for death | Model specific |
| A_q_ | 0.8 | – | ATP rate threshold for quiescence | Model specific |
| τ_min_ | 0.8 | day | Min cell cycle time | (10) |
| σ_min_ | 80 | μm | Min vessel spacing | (13) |
| σ_mean_ | 158 | μm | Mean vessel spacing | (13) |
| ν_mean_ | 20 | mM | Vessel stability parameter | Model specific |
| p_ang_ | 0.3 | – | Angiogenesis rate | Model specific |

**Table S6**

**Expression data used to fit linear models (Expression values are normalized counts from NanoString dataset)**

| **Phenotype** | **Gene** | **Expression** | **pH** | **a** | **b** | **c** | **d** | **e** |
| --- | --- | --- | --- | --- | --- | --- | --- | --- |
| M1 | *Fcer1g* | 9166.35 | 6.8 | -0.5 | -0.5 | 0 | 0 | -1 |
| M1 | *Fcer1g* | 9038.51 | 6.8 | -0.5 | -0.5 | 0 | 0 | -1 |
| M1 | *Fcer1g* | 8506.11 | 7.4 | -0.5 | -0.5 | 0 | 0 | -1 |
| M1 | *Fcer1g* | 8507.25 | 7.4 | -0.5 | -0.5 | 0 | 0 | -1 |
| M2 | *Fcer1g* | 4143.87 | 6.8 | 0 | 0 | 0.5 | 0.5 | 1 |
| M2 | *Fcer1g* | 3896.17 | 7.4 | 0 | 0 | 0.5 | 0.5 | 1 |
| M1 | *Cd206* | 868.25 | 6.8 | -0.5 | -0.5 | 0 | 0 | -1 |
| M1 | *Cd206* | 745.9 | 6.8 | -0.5 | -0.5 | 0 | 0 | -1 |
| M1 | *Cd206* | 600.33 | 7.4 | -0.5 | -0.5 | 0 | 0 | -1 |
| M1 | *Cd206* | 620.23 | 7.4 | -0.5 | -0.5 | 0 | 0 | -1 |
| M2 | *Cd206* | 9712.51 | 6.8 | 0 | 0 | 0.5 | 0.5 | 1 |
| M2 | *Cd206* | 6881.18 | 7.4 | 0 | 0 | 0.5 | 0.5 | 1 |
| M1 | *Tgfb1* | 2376.33 | 6.8 | -0.5 | -0.5 | 0 | 0 | -1 |
| M1 | *Tgfb1* | 2258.5 | 6.8 | -0.5 | -0.5 | 0 | 0 | -1 |
| M1 | *Tgfb1* | 2812.9 | 7.4 | -0.5 | -0.5 | 0 | 0 | -1 |
| M1 | *Tgfb1* | 2702.55 | 7.4 | -0.5 | -0.5 | 0 | 0 | -1 |
| M2 | *Tgfb1* | 3764.51 | 6.8 | 0 | 0 | 0.5 | 0.5 | 1 |
| M2 | *Tgfb1* | 3249.3 | 7.4 | 0 | 0 | 0.5 | 0.5 | 1 |
| M1 | *Tgfbr1* | 2024.85 | 6.8 | -0.5 | -0.5 | 0 | 0 | -1 |
| M1 | *Tgfbr1* | 2122.38 | 6.8 | -0.5 | -0.5 | 0 | 0 | -1 |
| M1 | *Tgfbr1* | 2933.21 | 7.4 | -0.5 | -0.5 | 0 | 0 | -1 |
| M1 | *Tgfbr1* | 2961.61 | 7.4 | -0.5 | -0.5 | 0 | 0 | -1 |
| M2 | *Tgfbr1* | 2789.48 | 6.8 | 0 | 0 | 0.5 | 0.5 | 1 |
| M2 | *Tgfbr1* | 3410.86 | 7.4 | 0 | 0 | 0.5 | 0.5 | 1 |
| M1 | *Ccl2* | 18274.44 | 6.8 | -0.5 | -0.5 | 0 | 0 | -1 |
| M1 | *Ccl2* | 18361.3 | 6.8 | -0.5 | -0.5 | 0 | 0 | -1 |
| M1 | *Ccl2* | 43221.27 | 7.4 | -0.5 | -0.5 | 0 | 0 | -1 |
| M1 | *Ccl2* | 42874.46 | 7.4 | -0.5 | -0.5 | 0 | 0 | -1 |
| M2 | *Ccl2* | 2818.95 | 6.8 | 0 | 0 | 0.5 | 0.5 | 1 |
| M2 | *Ccl2* | 4817.4 | 7.4 | 0 | 0 | 0.5 | 0.5 | 1 |
| M1 | *Ccrl2* | 2334.98 | 6.8 | -0.5 | -0.5 | 0 | 0 | -1 |
| M1 | *Ccrl2* | 2172.47 | 6.8 | -0.5 | -0.5 | 0 | 0 | -1 |
| M1 | *Ccrl2* | 2705.04 | 7.4 | -0.5 | -0.5 | 0 | 0 | -1 |
| M1 | *Ccrl2* | 2346.96 | 7.4 | -0.5 | -0.5 | 0 | 0 | -1 |
| M2 | *Ccrl2* | 43.72 | 6.8 | 0 | 0 | 0.5 | 0.5 | 1 |
| M2 | *Ccrl2* | 41.76 | 7.4 | 0 | 0 | 0.5 | 0.5 | 1 |

**Table S7**

**Coefficients for linear model fit to expression data**

| **Parameter** | **Value** | **Units** | | **Description** | **Reference** |
| --- | --- | --- | --- | --- | --- |
| α*_Ccl2_* | -140892.7783 | – | | *Ccl2* expression intercept | Fit to gene expression data |
| β*_Ccl2_* | 22273.70417 | – | | *Ccl2* pH coefficient | Fit to gene expression data |
| γ*_Ccl2_* | 121062.6283 | – | | *Ccl2* ecology coefficient | Fit to gene expression data |
| δ*_Ccl2_* | -18942.95417 | – | | *Ccl2* ecology-pH interaction coefficient | Fit to gene expression data |
| μ*_Ccl2_* | 0.0 | – | | Min *Ccl2* expression | Estimated from expression fit |
| ν*_Ccl2_* | 47169.53 | – | | Max *Ccl2* expression | Estimated from expression fit |
| α*_Ccrl2_* | -383.0625 | – | | *Ccrl2* expression intercept | Fit to gene expression data |
| β*_Ccrl2_* | 225.2625 | – | | *Ccrl2* pH coefficient | Fit to gene expression data |
| γ*_Ccrl2_* | 448.9958333 | – | | *Ccrl2* ecology coefficient | Fit to gene expression data |
| δ*_Ccrl2_* | -228.5291667 | – | | *Ccrl2* ecology-pH interaction coefficient | Fit to gene expression data |
| μ*_Ccrl2_* | 41.43333 | – | | Min *Ccrl2* expression | Estimated from expression fit |
| ν*_Ccrl2_* | 2571.37917 | – | | Max *Ccrl2* expression | Estimated from expression fit |
| α*_Tgfb1_* | 3465.395833 | – | | *Tgfb1* expression intercept | Fit to gene expression data |
| β*_Tgfb1_* | -62.41666667 | – | | *Tgfb1* pH coefficient | Fit to gene expression data |
| γ*_Tgfb1_* | 6138.160833 | – | | *Tgfb1* ecology coefficient | Fit to gene expression data |
| δ*_Tgfb1_* | -796.2666667 | – | | *Tgfb1* ecology-pH interaction coefficient | Fit to gene expression data |
| μ*_Tgfb1_* | 1950.490 | – | | Min *Tgfb1* expression | Estimated from expression fit |
| ν*_Tgfb1_* | 4193.852 | – | | Max *Tgfb1* expression | Estimated from expression fit |
| α*_Tgfbr1_* | -6041.110833 | – | | *Tgfbr1* expression intercept | Fit to gene expression data |
| β*_Tgfbr1_* | 1245.979167 | – | | *Tgfbr1* pH coefficient | Fit to gene expression data |
| γ*_Tgfbr1_* | 1788.284167 | – | | *Tgfbr1* ecology coefficient | Fit to gene expression data |
| δ*_Tgfbr1_* | -210.3458333 | – | | *Tgfbr1* ecology-pH interaction coefficient | Fit to gene expression data |
| μ*_Tgfbr1_* | 1345.452 | – | | Min *Tgfbr1* expression | Estimated from expression fit |
| ν*_Tgfbr1_* | 3514.423 | – | | Max *Tgfbr1* expression | Estimated from expression fit |
| α*_Fcer1g_* | 11402.7 | – | | *Fcer1g* expression intercept | Fit to gene expression data |
| β*_Fcer1g_* | -702.875 | – | | *Fcer1g* pH coefficient | Fit to gene expression data |
| γ*_Fcer1g_* | -4451.563333 | – | | *Fcer1g* ecology coefficient | Fit to gene expression data |
| δ*_Fcer1g_* | 290.0416667 | – | | *Fcer1g* ecology-pH interaction coefficient coefficient | Fit to gene expression data |
| μ*_Fcer1g_* | 3854.887 | – | | Min *Fcer1g* expression | Estimated from expression fit |
| ν*_Fcer1g_* | 9598.888 | – | | Max *Fcer1g* expression | Estimated from expression fit |
| α*_Cd206_* | 22419.1675 | – | | *Cd206* expression intercept | Fit to gene expression data |
| β*_Cd206_* | -2523.4375 | – | | *Cd206* pH coefficient | Fit to gene expression data |
| γ*_Cd206_* | 19381.74917 | – | | *Cd206* ecology coefficient | Fit to gene expression data |
| δ*_Cd206_* | -2195.445833 | – | | *Cd206* ecology-pH interaction coefficient | Fit to gene expression data |
| μ*_Cd206_* | 577.4808 | – | | Min *Cd206* expression | Estimated from expression fit |
| ν*_Cd206_* | 12071.9517 | – | | Max *Cd206* expression | Estimated from expression fit |
| D_CCL2_ | 30 | μm^2^/s  μm^2^/s | | Diffusion rate of *CCL2* |  |
| D _TGF-β_ | 26 | μm^2^/s  μm^2^/s | | Diffusion rate of TGF-β |  |
| λ | 0.99 | – | | Cytokine decay rate | Model specific |
| θ | 400 | – | | Strength of chemotaxis | Model specific |
| ψ | 200 | μm/day | | Maximum movement radius |  |
| ω_TGF-β_ | 800 |  | | TGF-β disassociation constant | Model specific |
| ω_CCL2_ | 10000 |  | | CCL2 disassociation constant | Model specific |
| N_M_ | 125 | cells | Number of macrophages | | Model specific |
| N_T_ | 25 | cells | Initial number of tumor cells | | Model specific |
