## Supplementary material for "Acidity promotes tumor progression by altering macrophage phenotype in prostate cancer"

**Supplemental figures and legends**

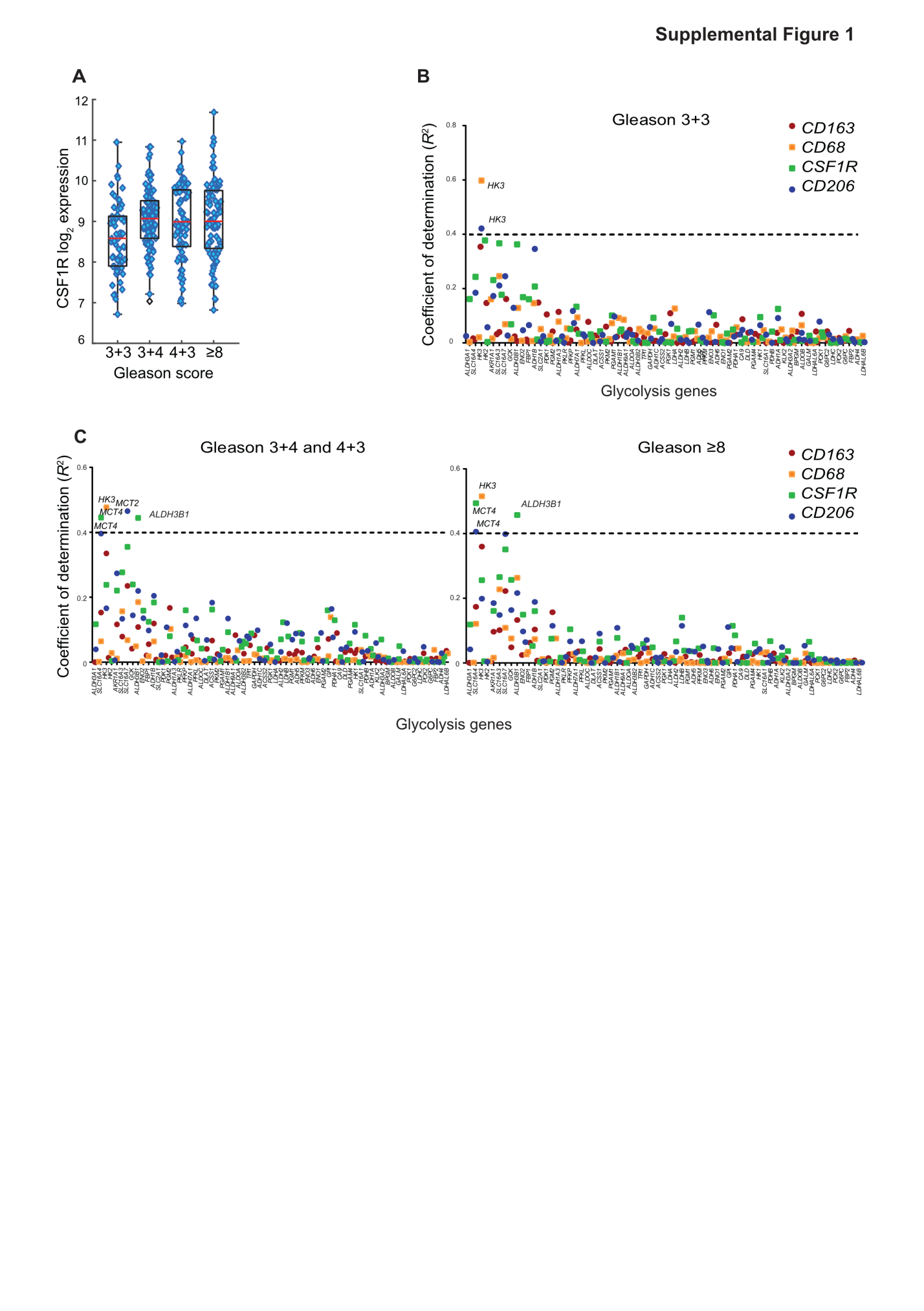
**Supplemental Figure 1: Macrophage infiltration correlates with MCT4 expression**

**A)** CSF1R expression in early stage patients (Gleason score= 3+3) and advanced prostate (Gleason score = 3+4, 4+3 and ≥8) retrieved from the TCGA PRAD333, p<0.05. **B-C)** plots of coefficient of determination (R^2^) between major macrophage markers and genes known to be regulating glycolysis (KEGG glycolysis pathway related genes, see Table S4) in patients with Gleason score 3+3 and Gleason score 4+3/4+3 or ≥8.

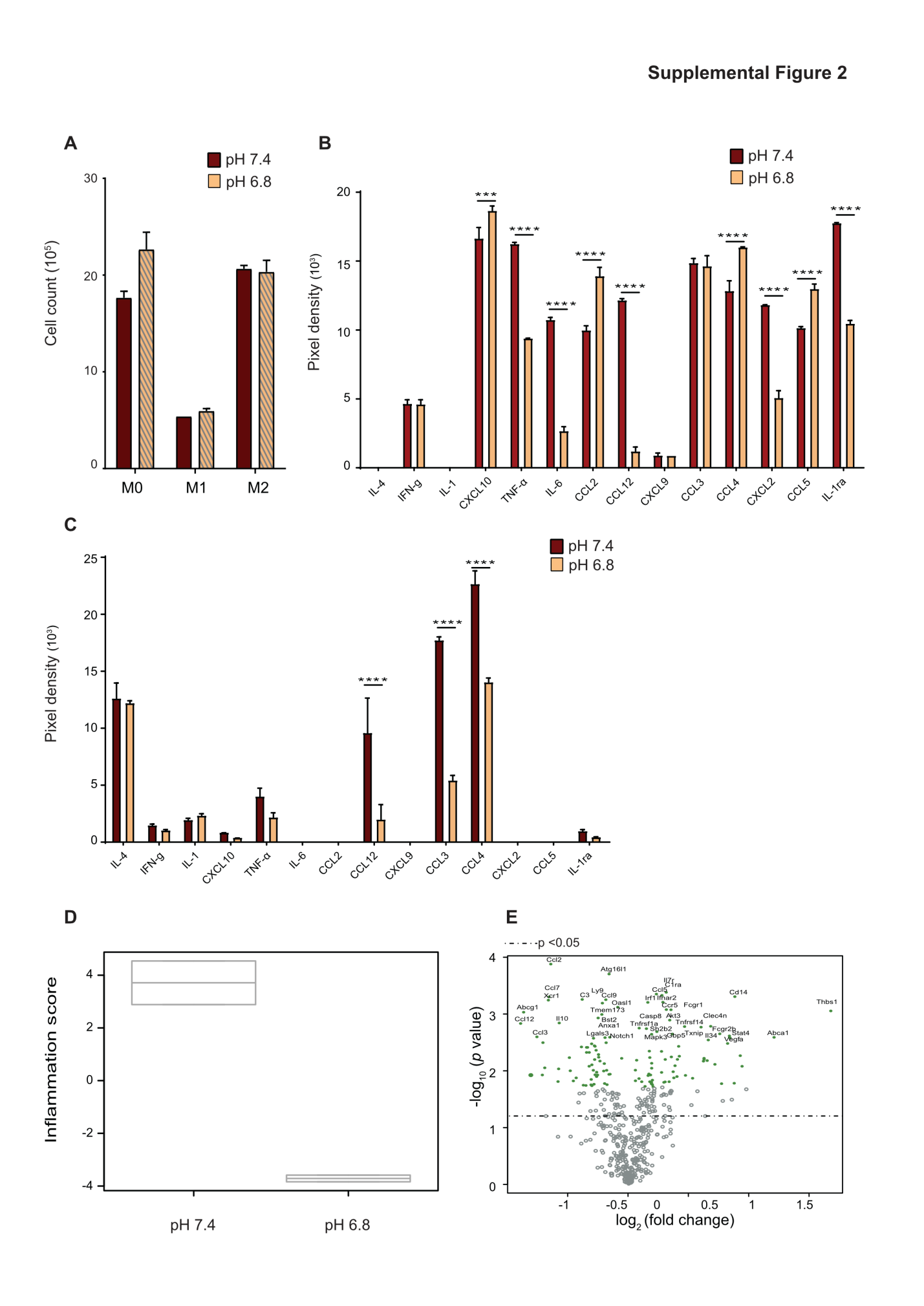
**Supplemental Figure 2: Extracellular acidosis alters macrophage activation *in vitro***

**A)** Cell count of BMDMs stimulated for 24 hr with LPS/IFN-γ (M1) or IL4 (M2), or left untreated as M0. **B-C)** Macrophages were stimulated at pH 7.4 or pH 6.8 using LPS/IFN-γ (B) or IL-4 (C). The culture medium of each phenotype was collected after 24 hr. Culture medium was then processed for determination of cytokines released using mouse cytokine array Panel A. Densitometric analysis was then done using Image J software and mean pixel density was graphed (n=2). **D-E)** Inflammation score (D) and Volcano plot (E) generated by nSolver software 3.0 using gene expression data from nCounter PanCancer Immune profiling of 6 h-LPS/IFN-γ stimulated macrophages activated at pH 7.4 compared with those activated at pH 6.8 (n=2).

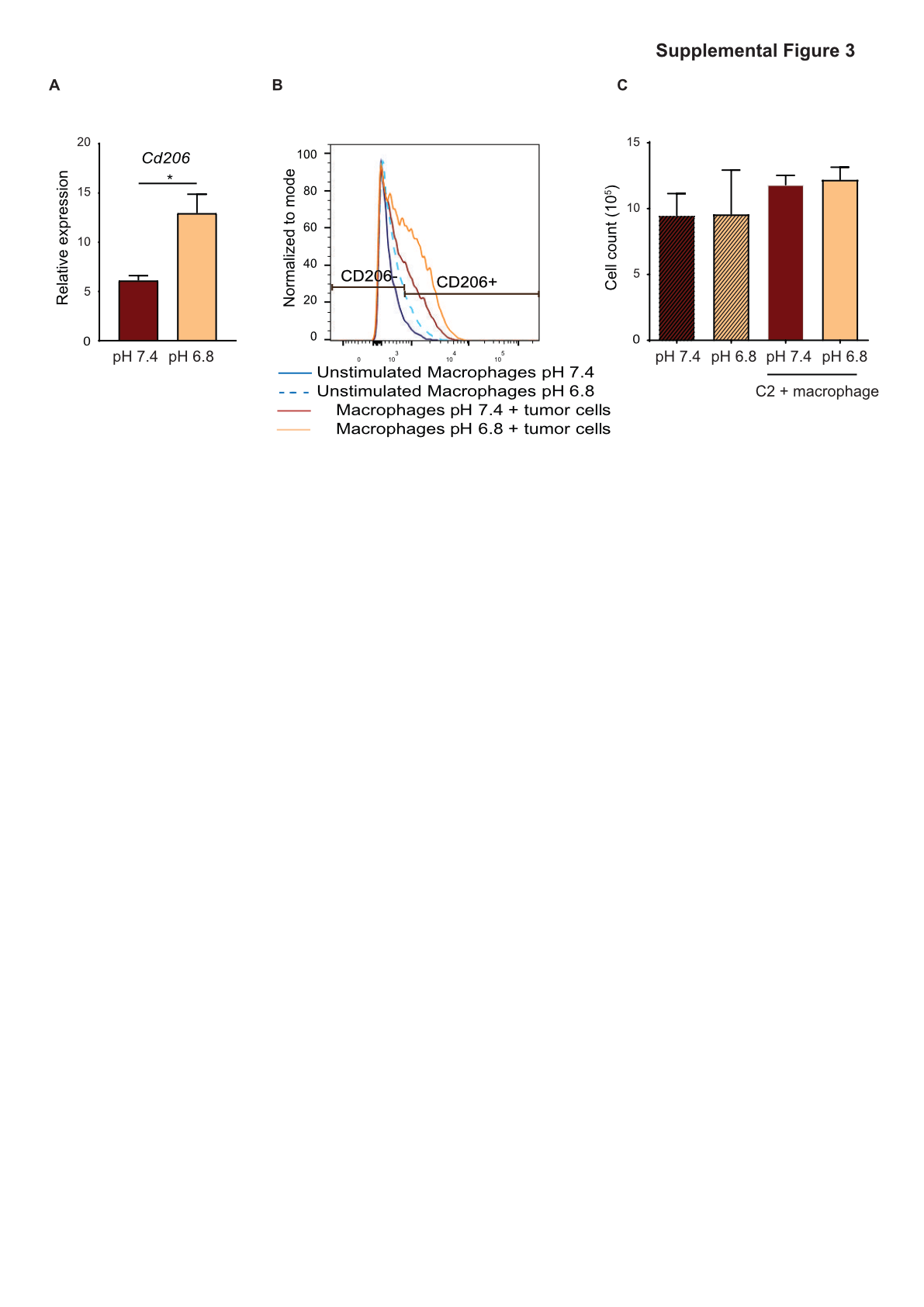

**Supplemental Figure 3: Extracellular acidosis enhances a tumor-promoting macrophage phenotype**

**A)** Relative mRNA level of *Cd206* in macrophages directly co-cultured with TRAMP-C3 for 6 days at pH 7.4 or pH 6.8, then sorted and processed for RNA extraction. **B)** Flow cytometry analysis of CD206 expression in macrophages incubated at pH 7.4 or pH 6.8 for 24 hr then either cultured alone or with TRAMP-C3 at pH 7.4 for another 24 hr. F4/80 staining was used to gate out tumor cells. **C)** Total cell count of TRAMP-C2 co-cultured with or without macrophages in neutral or acidic medium for 24 hr.

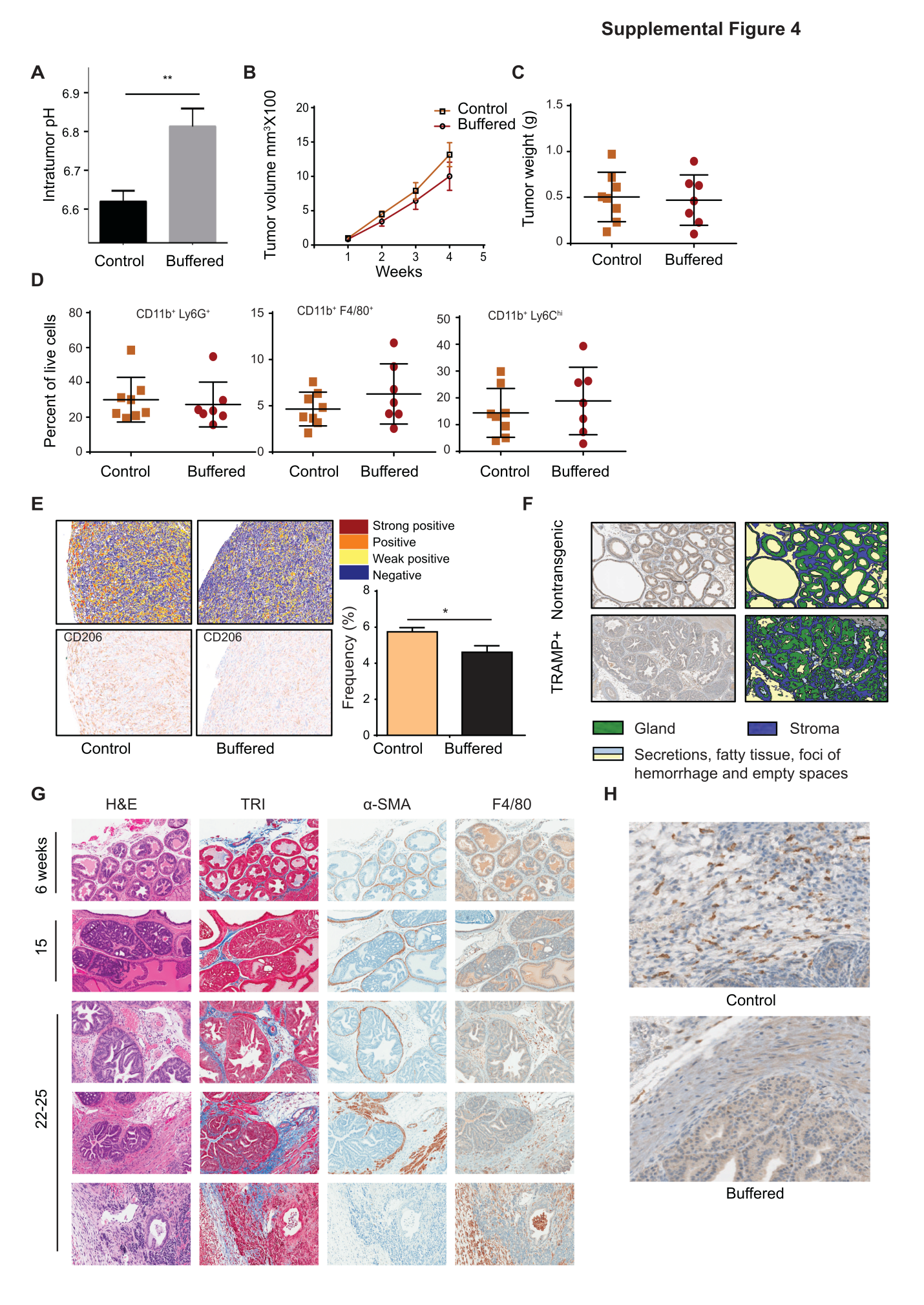

**Supplemental Figure 4: Buffering tumor-secreted acids alters TAM phenotype *in vivo* and reduces tumor progression**

**A)** Intratumoral pH measured *in vivo* by a microelectrode in control or sodium bicarbonate- treated (buffered) TRAMP-C2-bearing mice (n=7-8). **B)** Tumor volume monitored by digital caliper in control or sodium bicarbonate-treated (buffered) TRAMP-C2-bearing mice. **C)** Tumor weight. **D)** Single cell suspensions were processed for flow cytometric staining using antibodies cocktail specific for CD11b, Ly6G, Ly6C and F4/80. Ly6G and Ly6C staining was then used to identify CD11b^+^ Ly6G^+^Ly6C ^low^ (granulocytic MDSCs), CD11b^+^Ly6C^hi^Ly6G^-^ (monocytic MDSCs) and CD11b^+^Ly6G^-^Ly6C^-^F4/80^+^ (macrophages) subsets. **E**) Representative images of CD206-stained tumor sections **F)** Representative images of tissue segmentation performed using Definiens Tissue Studio v4.0 suite. **G)** Representative images of serial sections of paraffin embedded prostates, isolated from 6, 15, 22, 23 and 25 weeks old TRAMP mice, stained using H&E, Masson’s Trichome (TRI, collagen), α-SMA (fibroblast) and F4/80 (macrophage). **H)** Representative images of tumor sections stained with F4/80 collected from of 32 weeks old TRAMP mice treated with 200 mM sodium bicarbonate in their drinking water starting from 4 weeks of age (buffered) or kept on tap water as control.

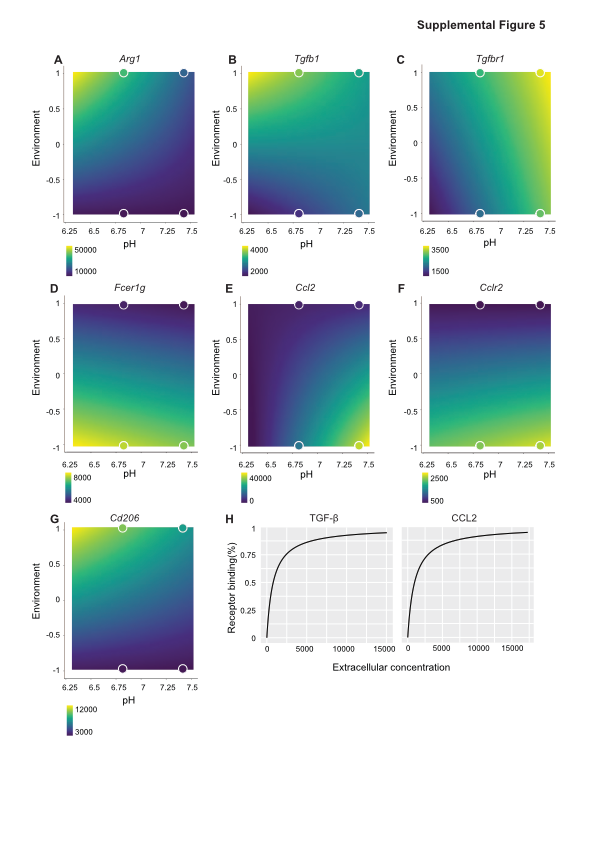

**Supplemental Figure 5: Acid-responsive macrophages promote tumor growth *in silico* A-G)** Output of linear model fitting of *Arg1, Tgfb1, Tgfbr1, Fcer1g, Ccl2, Cclr2* and *Cd206* expression represented as heatmap. For each phenotypic trait, a linear model allows to predict expression under a variety of conditions. Here, -1 is a tumor rich inflammatory environment, while 1 is environment with necrosis and anti-inflammatory cytokines. The circles outlined in white are the actual *in vitro* data. **H)** Binding of TGF-β by TGF-βR, and CCL2 by CCLR2 was modeled using the Hill equation.

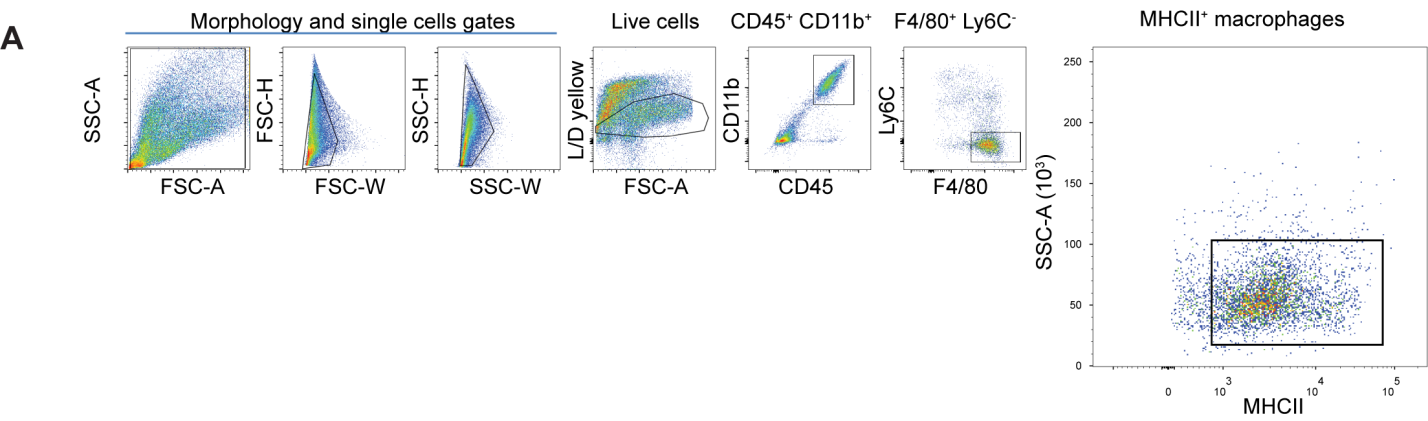

**Supplemental Figure 6:**

**A)** Gating strategy used to sort TAMs: single cell suspensions were incubated with live/dead (L/D) fixable yellow fluorescent reactive dye to exclude dead cells then stained with conjugated antibodies against surface markers; CD45, CD11b, Ly6C, F4/80 and MHCII.
